# Spatiotemporal dynamics of cytokines expression dictate fetal liver hematopoiesis

**DOI:** 10.1101/2023.08.24.554612

**Authors:** Marcia Mesquita Peixoto, Francisca Soares-da-Silva, Valentin Bonnet, Gustave Ronteix, Rita Faria Santos, Marie-Pierre Mailhe, Xing Feng, João Pedro Pereira, Emanuele Azzoni, Giorgio Anselmi, Marella de Bruijn, Charles N. Baroud, Perpétua Pinto-do-Ó, Ana Cumano

## Abstract

During embryogenesis, yolk-sac and intra-embryonic-derived hematopoietic progenitors, comprising the precursors of adult hematopoietic stem cells, converge into the fetal liver. With a new staining strategy, we defined all non-hematopoietic components of the fetal liver and found that hepatoblasts are the major producers of hematopoietic growth factors. We identified mesothelial cells, a novel component of the stromal compartment, producing Kit ligand, a major hematopoietic cytokine.

A high-definition imaging dataset analyzed using a deep-learning based pipeline allowed the unambiguous identification of hematopoietic and stromal populations, and enabled determining a neighboring network composition, at the single cell resolution.

Throughout active hematopoiesis, progenitors preferentially associate with hepatoblasts, but not with stellate or endothelial cells. We found that, unlike yolk sac-derived progenitors, intra-embryonic progenitors respond to a chemokine gradient created by CXCL12-producing stellate cells. These results revealed that FL hematopoiesis is a spatiotemporal dynamic process, defined by an environment characterized by low cytokine concentrations.

## Introduction

The generation of hematopoietic cells occurs in different embryonic locations at successive, overlapping, time windows^(1,2)^. In the mouse, multipotent erythro-myeloid progenitors (EMP)^(3)^ are produced through an endothelial-to-hematopoietic transition (EHT) process, in the vasculature of the YS^(4)^, between E8-8.5. These progenitors differentiate into definitive erythrocytes^(5)^, tissue resident macrophages that can persist for life^(6)^, neutrophils and mast cells^(7,8)^ but are devoid of lymphoid potential^(3,9)^. Starting at E9, a subset of endothelial cells in the intra-embryonic (IE) vasculature (dorsal aorta, omphalo-mesenteric and vitelline arteries) undergoes EHT, spanning for around 48 hours and resulting in the emergence of the first lympho-myeloid progenitors^(10–15)^. Newly generated multipotent progenitors rapidly enter the circulation and home to the FL, where all hematopoietic generations converge. It is in this environment that the multipotent progenitors give rise to the hematopoietic stem cell (HSC) compartment. To acquire long-term reconstitution (LTR) capacity, IE multipotent progenitors require maturation that can occur in the dorsal aorta but primarily takes place in the FL^(16–18)^. Therefore, only few LTR HSC have been detected in E11.5 dorsal aorta and are only consistently found in the FL at around E12.5^(11,19,20)^. KIT is expressed from very early stages but other adult markers of HSC, such as Sca-1 and CD150, are only clearly detectable at E12.5 and E13.5, respectively^(18)^. The most substantial increase in FL reconstitution capacity occurs between E12.5-E16.5^(20)^. The early stages after FL colonization (E12.5-E14.5) constitute a unique environment where the HSC compartment is established.

The FL is an organ composed of hepatoblasts, of endodermal origin, that later differentiate into hepatocytes and cholangiocytes, being the latter dependent on signals from mesenchymal cells around the portal vessels, for differentiation^(21,22)^. Mesothelial cells that form the liver capsule derive from the septum transversum and can generate fibroblasts through epithelial-to-mesenchymal transition (EMT)^(21)^. Stellate cells that mature into vitamin A storing cells, can be found adjacent to mesothelial cells and lining the sinusoids^(23,24)^. An additional population of perivascular fibroblasts identified as NG2^+^, are found exclusively around portal vessels^(25)^. After E16.5 the liver transits from a hematopoietic to a metabolic organ and hepatoblasts rapidly differentiate into hepatocytes and cholangiocytes.

Several studies identified stromal compartments involved in fetal hematopoiesis. DLK1 expressing cells (defining hepatoblasts) were reported as main producers of erythropoietin (Epo) but also of stem cell factor (also called Kit-ligand (KITL)), and CXCL12^(26)^. Endothelial^(27,28)^ and stellate cells have been found to produce KITL^(29)^ and peri-portal NG2^+^ cells producing CXCL12 postulated as essential for HSC maintenance^(25)^. However, deletion of NG2 expressing peri-portal cells^(25)^ and constitutive deletion of KITL in hepatoblasts resulted in subtle reductions, or no changes, in the HSC compartment before birth^(29)^. In addition, most studies of the FL stroma restricted their analysis to particular subsets, concentrated in single time-points and did not consider the early stages of hematopoiesis.

Here we optimized 2D-3D imaging to the analysis of all stromal cell populations. The high-quality images were analyzed in a pipeline based in deep learning to determine the identity of individual cells and to describe their immediate neighbors. We show that hematopoietic progenitors of YS and IE have a higher frequency at the vicinity of the FL sub-mesothelial region, dominated by CXCL12-producing stellate cells and KITL-producing hepatoblasts and mesothelial cells. Later in development, mesenchymal cells (mainly perivascular) increase CXCL12 expression and CXCR4-expressing IE progenitors, but not CXCR4^−^ YS-progenitors, distribute away from the sub-mesothelial region. We identified mesothelial cells as producers of KITL and stellate cells/pericytes of CXCL12. Hepatoblasts, however, are major producers of KITL, CXCL12, and the only subset that produces Epo and interleukin7 (IL-7). Through E12.5 and E14.5 hematopoietic progenitors preferentially associate with hepatoblasts, but not with stellate or endothelial cells. By E18.5 hepatic cells that now consist of a majority of hepatocytes and cholangiocytes produce decreased levels of KITL, CXCL12, Epo and IL-7. This is coincident with the onset of BM hematopoiesis. Throughout FL development cytokine and chemokine production is strikingly lower compared with BM derived stromal cells, a feature that might be determinant for FL hematopoiesis.

## Results

### The non-hematopoietic compartment of the FL comprises endothelial, hepatic, mesenchymal and mesothelial cells

We integrated spectral flow cytometry and single-cell transcriptional profiling of isolated FL cells with an imaging pipeline capable of spatial analysis at the cellular level (Fig 1A). Using a set of defined surface markers (TableS1) we characterized the hematopoietic compartment at E12.5 (Fig1B), which predominantly consisted of erythroblasts (80% of total FL cells), a population of CD45^−^ erythroid progenitors (P1, P2, P3) that we previously showed derive from the YS (10%)^(5)^, and CD45^+^ IE-derived cells (5%) (Fig1B; FigS1A). These three major components of the hematopoietic compartment can be identified in the tissue with the combination of the erythroid markers Ter119 and CD71, the pan-hematopoietic marker CD45 and the progenitor marker KIT (Fig1C). A combination of CD45 and KIT staining allows the identification of three major non-erythroid populations in the FL. These include CD45^+^KIT^−^ cells, referred to as CD45^+^ cells, which comprise monocytes and granulocytes; CD45^−^KIT^+^ cells, referred to as KIT^+^ progenitors, that comprise the YS-derived erythroid progenitors^(5)^; and CD45^+^KIT^+^, referred to as DP (Double Positive) progenitors, that include a majority of common myeloid progenitors (CMP), granulocyte-monocyte progenitors (GMP), some of which can be of YS origin at E12.5 (not more than 8%^(30)^), megakaryocyte-erythroid progenitors (MEP), as well as LSK CD48^−^ and LSK CD48^+^ multipotent progenitors (FigS1B-D).

**Figure 1.**
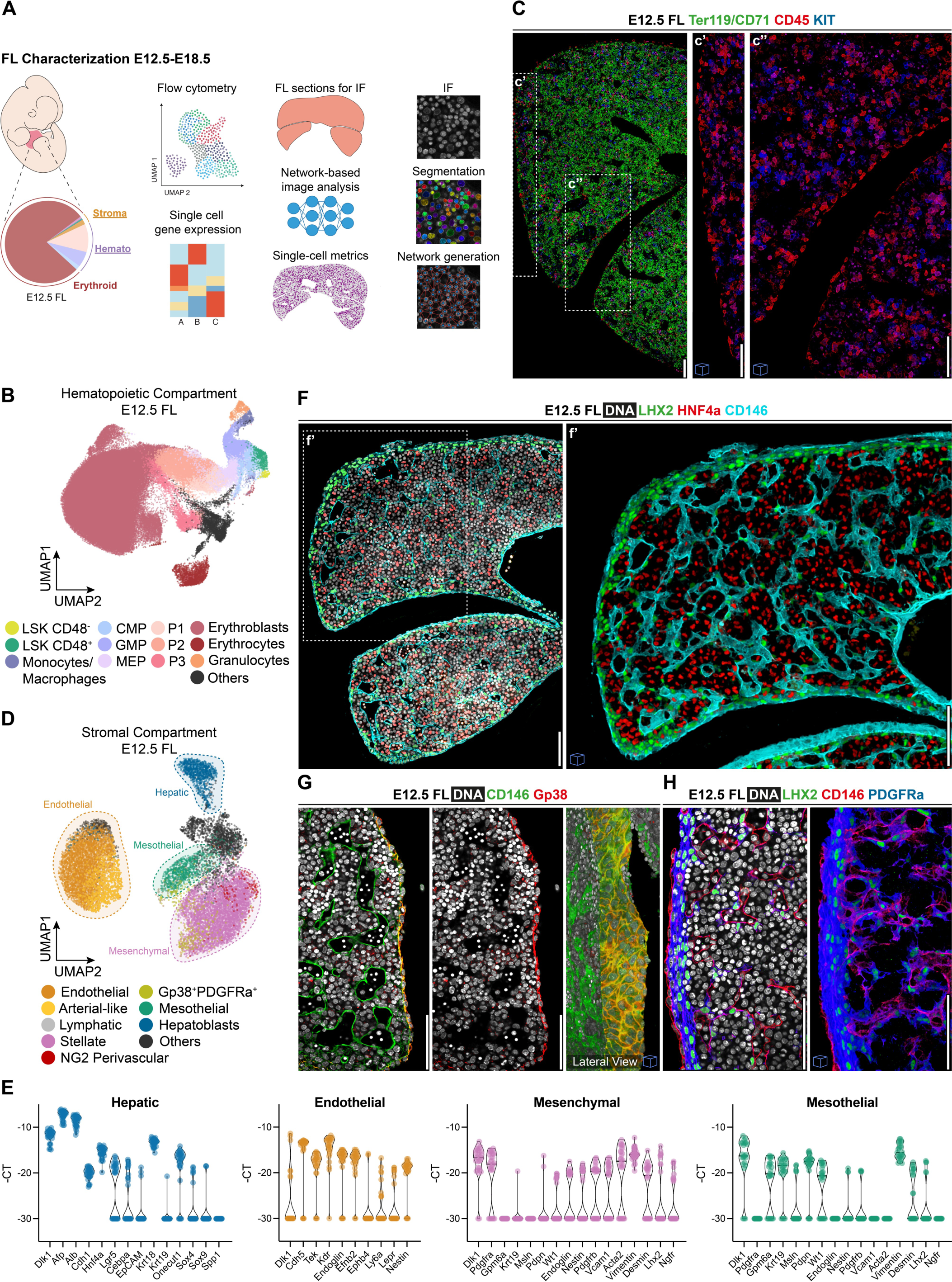
The non-hematopoietic compartment of the FL comprises endothelial, hepatic, mesenchymal and mesothelial cells. **(A)** Schematic representation of the strategy for the analysis of the FL from E12.5 to E18.5. **(B)** UMAP analysis of flow cytometry data of E12.5 FLs stained with the surface markers Ter119, CD45, CD71, Gr1, CD11b, KIT, Sca1, CD16/32, CD34, CD150 and CD48. See gating strategy used in Fig S1A. **(C)** Single-stack immunohistochemistry (IHC) of E12.5 FL with the hematopoietic markers Ter119&CD71 (green), CD45 (red) and KIT (blue). Inserts c’ and c" show an enlarged 3D view (20 um) of the selected regions with the markers CD45 and KIT. Scale bar 50 um. **(D)** UMAP analysis of flow cytometry data of Ter119^−^CD45^−^CD71^−^KIT^−^ cells from E12.5 FLs stained with the surface markers CD146, CD31, Gp38, PDGFRa, NG2, CD166, Sca1, E-Cadh and EpCAM that identify the four major populations highlighted by dashed circles: endothelial, hepatic, mesothelial and mesenchymal. See gating strategy used in Fig S2A. **(E)** Violin plots of single-cell gene expression analysis of lineage associated transcripts of sorted hepatic (CD31^−^Gp38^−^PDGFRa^−^EpCAM-E-Cadh^hi^ cells, blue), endothelial (CD31^+^CD146^+^ cells, orange), mesenchymal (CD31^−^Gp38^−^PDGFRa^+^ cells, pink), and mesothelial (CD31^−^PDGFRa^−^Gp38^+^ cells, green) cells. **(F)** Single-stack IHC of E12.5 FL with DAPI (white), LHX2 (green), the hepatic marker HNF4a (red) and CD146 (cyan). Insert f’ shows an enlarged 3D view (15 um) of the selected region. Scale bar 50 um. **(G)** Single-stack IHC of the capsular region of E12.5 FL with DAPI (white), the endothelial and mesothelial marker CD146 (green) and the mesothelial marker Gp38 (red) (left), Gp38 alone (middle), and a 3D lateral view (50 um) of both CD146 and Gp38 (right). Scale bar 50 um. **(H)** Single-stack IHC of the border region of E12.5 FL with DAPI (white), CD146 (red), the mesenchymal markers LHX2 (green) and PDGFRa (blue) (left), and a 3D view (20 um) (right). Scale bar 50 um. All images with cubes are 3D projections.

We performed spectral flow cytometry analysis of the refined non-hematopoietic compartment (Ter119^−^CD45^−^CD117^−^CD71^−^) with a panel of 14 surface markers (TableS1) (FigS2A-C). We identified 10 cell types that could be classified into 4 major populations: endothelial (CD31^+^CD146^+^), hepatic (CD31^−^Gp38^−^PDGFRa^−^E-Cadh^+^), mesenchymal (CD31^−^Gp38^−^PDGFRa^+^), and mesothelial (CD31^−^ PDGFRa^−^Gp38^+^) cells (Fig1D, FigS2A-C).

**Figure 2.**
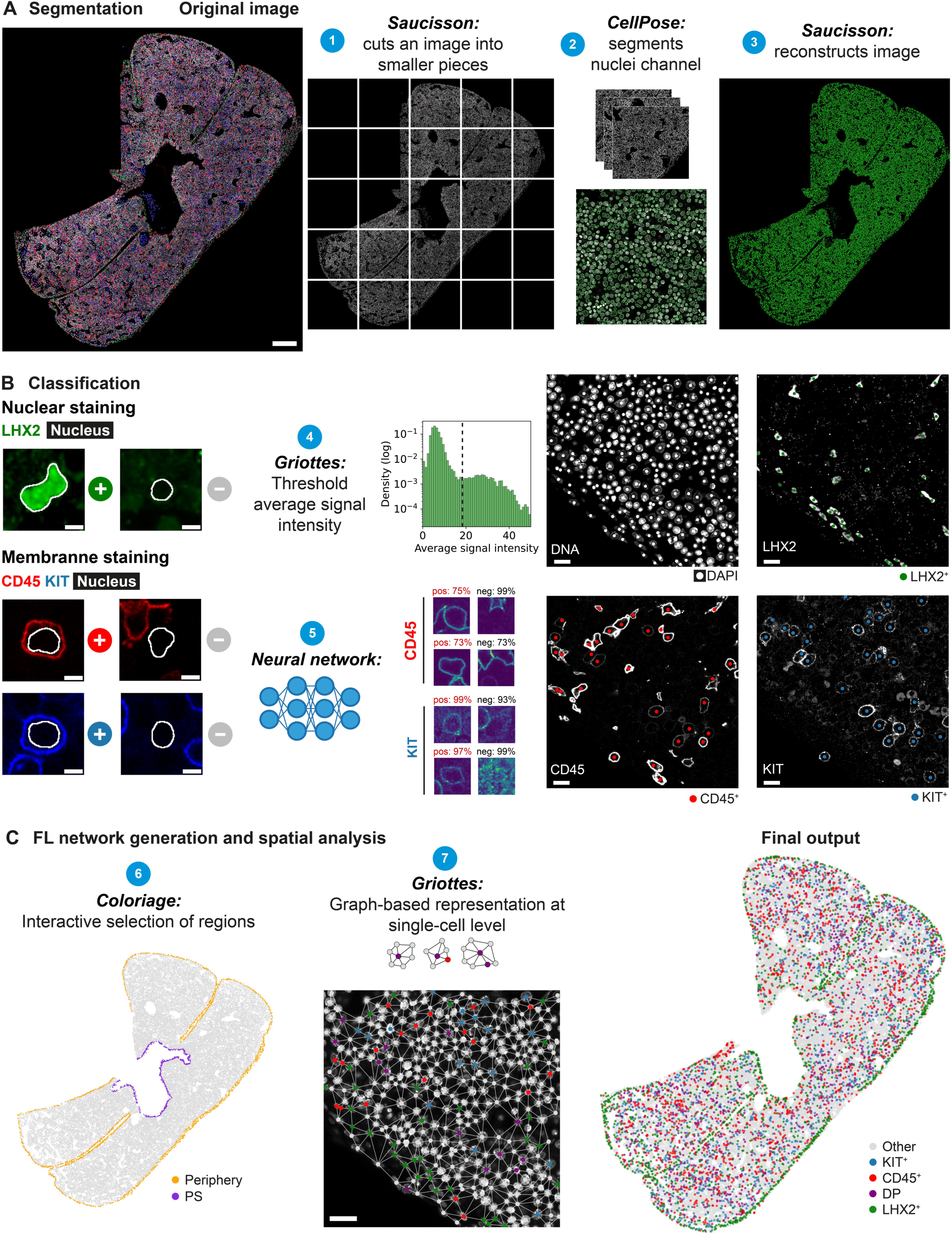
Image analysis workflow. **(A)** Segmentation pipeline. FL multi-channel images are cut into small pieces using *Saucisson* (1), Individual tiles are segmented based on DAPI signal using *CellPose* (2), and then reassembled into the original shape (3). On the left, a representative single-stack IHC of E12.5 FL with DAPI (white), LHX2 (green), KIT (blue) and CD45 (red) is shown to demonstrate the pipeline. Scale bar 200 μm. **(B)** Cell type classification pipeline. In case of nuclear staining (as for LHX2, green), cells are classified according to a threshold set on the average signal intensity inside the nuclei mask using *Griottes* (4). For membrane staining (as for CD45/red, KIT/blue), cells are classified using a *Neural Network* (5). A representative example of a small tile post-classification is presented, displaying positive cells for LHX2 (green), CD45 (red) and KIT (blue). Scale bar 5 μm (small tiles on the left), 10 μm (on the tiles on the right). **(C)** For FL network generation and spatial analysis, *Coloriage* is used to remove or highlight specific regions of the FL sections (6). *Griottes* is used to represent the cells in a dot plot and to plot the contacts between cells (7). Scale bar 10 μm. PV – *portal sinus*.

During the early stages of development (E12.5), E-Cadh^+^ hepatic cells consist primarily of E-Cadh^hi^EpCAM^−^ hepatoblasts (FigS2A-C) that express high levels of *Dlk1*, *Afp*, *Alb*, *Vcam1*, *Alcam* and *Hnf4a*, as well as the progenitor marker *Lgr5* and the proliferation marker *Ki-67* (Fig1E; FigS2D, Cluster IV) and are identified in the tissue by their expression of HNF4a (Fig1F). At E18.5, E-Cadh^int^ cells were classified as hepatocytes because of lower expression levels of *Dlk1*, *Afp*, *Igf2*, *Vcam1*, the absence of *Lgr5* and *Vimentin* expression (FigS2A and S2D, Cluster II) and by the expression of the mature marker *Cebpa*. Starting from E16.5, E-Cadh^hi^EpCAM^+^ biliary duct cells or cholangiocytes, which express *Sox4*, *Sox9*, and *Spp1* but not *Dlk1* (FigS2A, D, Cluster I) can be identified in the tissue exclusively close to NG2^+^ cells, around portal vessels (FigS2G, H).

Throughout all stages analyzed, CD31^+^ endothelial cells expressed *Cdh5*, *Tek*, *Kdr*, *Endoglin* and *Efnb2*, and intermediate levels of *Nestin* and *Lepr* (Fig1E, FigS2E). We identified an endothelial subset co-expressing Sca-1, and another, co-expressing Gp38 (Podoplanin), associated with lymphatic vessels^(31)^ (FigS2A-C). Gp38^+^ endothelial cells clustered separately and were characterized by expression of *Dlk1*, *Wt1*, *Alcam*, *Krt18* and *Krt19* and lower levels of *Endoglin*, *Cdh5* and *Tek* (Cluster II, FigS2E). In the tissue, endothelial cells were detected by the expression of CD146, (Fig1F-H) that also labeled the mesothelial layer, identified by Gp38 co-expression (Fig1G).

PDGFRa^+^CD166^+^NG2low/^−^ fibroblasts represented an enriched population of maturing stellate cells co-expressing *Lhx2* and *Desmin* that acquire a specific autofluorescent profile as they accumulate vitamin A^(24)^ (FigS2F, Cluster I, II). In contrast, PDGFRa^+^CD166^−^NG2^+^ perivascular fibroblasts do not accumulate vitamin A granules, were barely detectable at E12.5 and increased in frequency as development progressed. We found that NG2^+^ cells and stellate cells exhibited similar transcriptional profiles based on the expression of *Endoglin*, *Nestin*, *Pdgfrb*, and *Vcam1* and clustered together (FigS2F, Cluster I, II). However, stellate cells, that co-express PDGFRa and LHX2 or Desmin, were located in the sub-mesothelial region and lining CD146^+^ endothelial cells (FigS2I) whereas NG2^+^ fibroblasts were exclusively found lining portal vessels, as previously described^(25)^ (Fig1F-G; FigS2H).

Mesothelial cells expressed *Gpm6a*, *Msln*, and *Wt1*, but stellate markers (Fig1H; FigS2F, Cluster III). A small subset of cells co-expressed PDGFRa, Gp38 and NG2, but not CD166 and co-expressed transcripts of both mesothelial and mesenchymal cells, consistent with a mesothelial origin of stellate cells^(32)^ (FigS2F, Cluster III). In the tissue, mesothelial cells formed a single epithelial sheet followed by an irregular layer of sub-mesothelial PDGFRa^+^LHX2^+^ cells, that can reach a thickness of 3-5 cells at E12.5. As development progresses, the sub-mesothelial region was consistently found to be a single cell layer, due to the relocation of stellate cells to a perivascular position (FigS2I). Sub-mesothelial and the peri-vascular stellate cells were only distinguishable by position in the tissue. Interestingly, both mesenchymal and mesothelial cells expressed the *Dlk1* transcript, which has been used as a marker for the identification and isolation of hepatoblasts^(26)^ (Fig1H).

Overall, three markers are sufficient to identify the stromal compartment. LHX2^+^ stellate cells that localize in the sub-mesothelium and around blood vessels, HNF4a^+^ hepatoblasts that distribute throughout the parenchyma and CD146^+^ endothelial and mesothelial cells that are distinguishable by their specific locations (Fig1F).

### YS and IE-derived progenitors accumulate at the border at E12.5 and are preferentially in contact with hepatoblasts, but not stellate or endothelial cells

The spatial distribution of different cell types was quantified by building a multi-step image analysis pipeline, that we named Pyxidis, to identify the phenotype and position of each cell within 2D and 3D images (Fig2, FigS3). The output of this pipeline was a graph-based representation of the cells that preserved spatial positioning and cell identity in a compact format^(33)^, thus enabling information from many different images to be pooled together for statistical analysis. Details of the analysis are described in materials and methods.

Classification of endothelial cells using a nuclear marker was more accurate than with surface CD146, therefore we analyzed FLs from the FLK1-GFP mouse reporter line (FigS4A). To validate the cell classification, we compared the frequencies of classified cells in the tissue with the frequencies obtained by flow cytometry analysis. Cells from the hematopoietic compartment were found with similar frequencies in the two methods (FigS4B). Stromal cells, in contrast, were over-represented by a factor of 3 in the imaging analysis due to constraints in the single cell isolation procedure (FigS4C). Overall, the image analysis pipeline provided a robust method for the study of different cell types in the tissue.

The graph-based description of the FL provided a global view of the spatial distribution of the cell types. CD45^+^ cells were spread in the parenchyma, while YS-derived KIT^+^ and IE-derived DP progenitors displayed a biased distribution towards the periphery, that we further quantified (Fig3A, C, E). For that, a random distribution of cells was computed (See Methods for detailed description) and compared to the frequency of a given cell type at a certain position (% observed/random). CD45^+^ cells had a random distribution. Conversely, KIT^+^ and DP progenitors were found close to the capsule (Fig 3E). Mesenchymal cells accumulate in the sub-mesothelial region (within 30 µm of the border) or distributed throughout the parenchyma, lining the vasculature, as mentioned above (Fig3B, D, F). Endothelial cells were evenly distributed throughout the parenchyma (Fig3F). Interestingly, HNF4a^+^ hepatoblasts displayed a similar distribution profile as hematopoietic progenitors with significantly higher frequencies near the sub-mesothelial region (Fig3F). In contrast, when analyzing the distribution of these populations in relation to a central region, the portal sinus (PS), we found that CD45^+^ cells were also randomly distributed, while KIT^+^ and DP progenitors were located further away (FigS4E). LHX2^+^ cells lined this vessel indicating that LHX2 is also expressed around non-sinusoidal vasculature (FigS4E).

**Figure 3.**
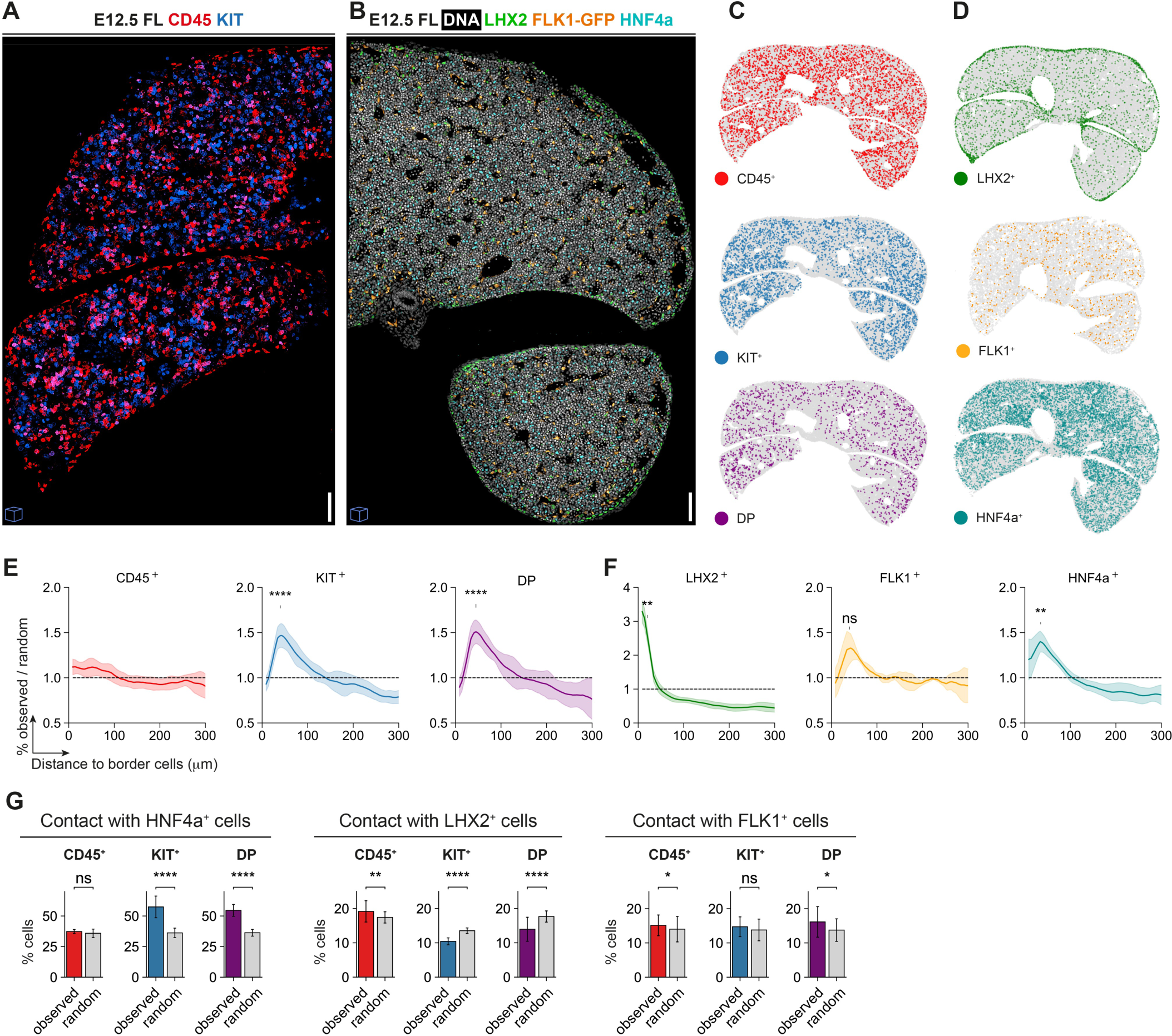
YS and IE-derived progenitors accumulate at the border at E12.5 and are preferentially in contact with hepatoblasts, but not stellate or endothelial cells. **(A)** IHC of 2 lobes of E12.5 FL with DAPI (white), CD45 (red) and KIT (blue), 3D view (15 μm). **(B)** IHC of 2 lobes of E12.5 FL (FLK1-GFP mice) with DAPI (white), LHX2 (green), FLK-1 GFP (orange), and HNF4a (cyan), 3D view (10 μm). **(C)** Dot plot representation of CD45^+^ (red), KIT^+^ (blue), and DP (purple) cells distribution of E12.5 FL representative sections (15 μm section projected into a single plane). **(D)** Dot plot representation of LHX2^+^ (green), FLK1^+^ (orange), and HNF4a^+^ (cyan) cells distribution of E12.5 FL representative sections (15 μm section projected into a single plane, except for the FLK1, that corresponds to a single-stack). **(E)** Distance to the capsule profile of CD45^+^ (red, n=10), KIT^+^ (blue, n=10), and DP (purple, n=10) cells of E12.5 FLs. Curves indicate average ± standard deviation. Statistical analysis was calculated at the peak of each curve using the Mann-Whitney test, between observed and random curves. *, P < 0.05; **, P < 0.01; ****, P < 0.0001; ns, not significant. **(F)** Distance to the capsule profile of LHX2^+^ (green, n=6), FLK1^+^ (orange, n=3), and HNF4a^+^ (cyan, n=6) cells of E12.5 FLs. Curves indicate average ± standard deviation. Statistical analysis was calculated at the peak of each curve using the Mann-Whitney test, between observed and random curves. *, P < 0.05; **, P < 0.01; ****, P < 0.0001; ns, not significant. **(G)** Neighborhood composition of CD45^+^, KIT^+^ and DP cells of E12.5 FLs. Graphs display the percentage of cells that are in contact with HNF4a^+^ (left, n=3), LHX2^+^ (middle, n=3) and FLK1^+^ (right, n=2) stromal cells (observed) compared to random. Bar plots indicate average ± standard deviation. Statistical analysis was calculated using t-test Welch. *, P < 0.05; **, P < 0.01; ****, P < 0.0001; ns, not significant. Images with cubes are 3D projections.

Analysis of cell-cell contacts showed that KIT^+^ and DP progenitors exhibited a preferential contact with hepatoblasts while they were found to be significantly less in contact with stellate cells (Fig3G). Unlike previous reports, we failed to find any preferential proximity of hematopoietic progenitors with FLK1^+^ endothelial cells at E12.5^(25,29)^ (Fig3G). Our results identify the sub-capsular region as a privileged site for hematopoietic progenitors of YS and IE origin at E12.5, that preferentially associate with hepatoblasts.

### Supportive hematopoietic cytokines are produced by distinct stromal populations and accumulate at the mesothelial and sub-mesothelial regions

We analyzed cytokine expression in single cells of the four major stromal populations and found that hepatoblasts were the major cytokine producers at E12.5 (Fig4A) expressing high levels of *Cxcl12* and *Kitl*, intermediate levels of *Csf1* and were the only producers of *Epo* and *Il7*. Mesothelial and endothelial cells were also major producers of *Kitl* and expressed intermediate levels of *Csf1*. *Cxcl12* was produced at high levels by virtually all mesenchymal cells (Fig4A, FigS2, Cluster I and II). Hepatoblasts that co-expressed *Epo* and *Il7* in addition to *Cxcl12* and *Kitl*, clustered together (FigS2D, Cluster IV). They were also the most immature cells because they co-expressed *Lgr5* and *Ki67*. In contrast, *Csf1* and *Cxcl12* were co-expressed by different cells within cluster IV and in E18.5 cluster II. *Csf1* expressing mesenchymal cells were mostly represented in Cluster II, that differ from Cluster I by the expression of the stellate markers *Lhx2* and *Ngfr* (FigS2F). Interestingly, *Igf2*, a crucial embryonic growth factor, was highly expressed in all analyzed stromal populations (FigS2D-F).

**Figure 4.**
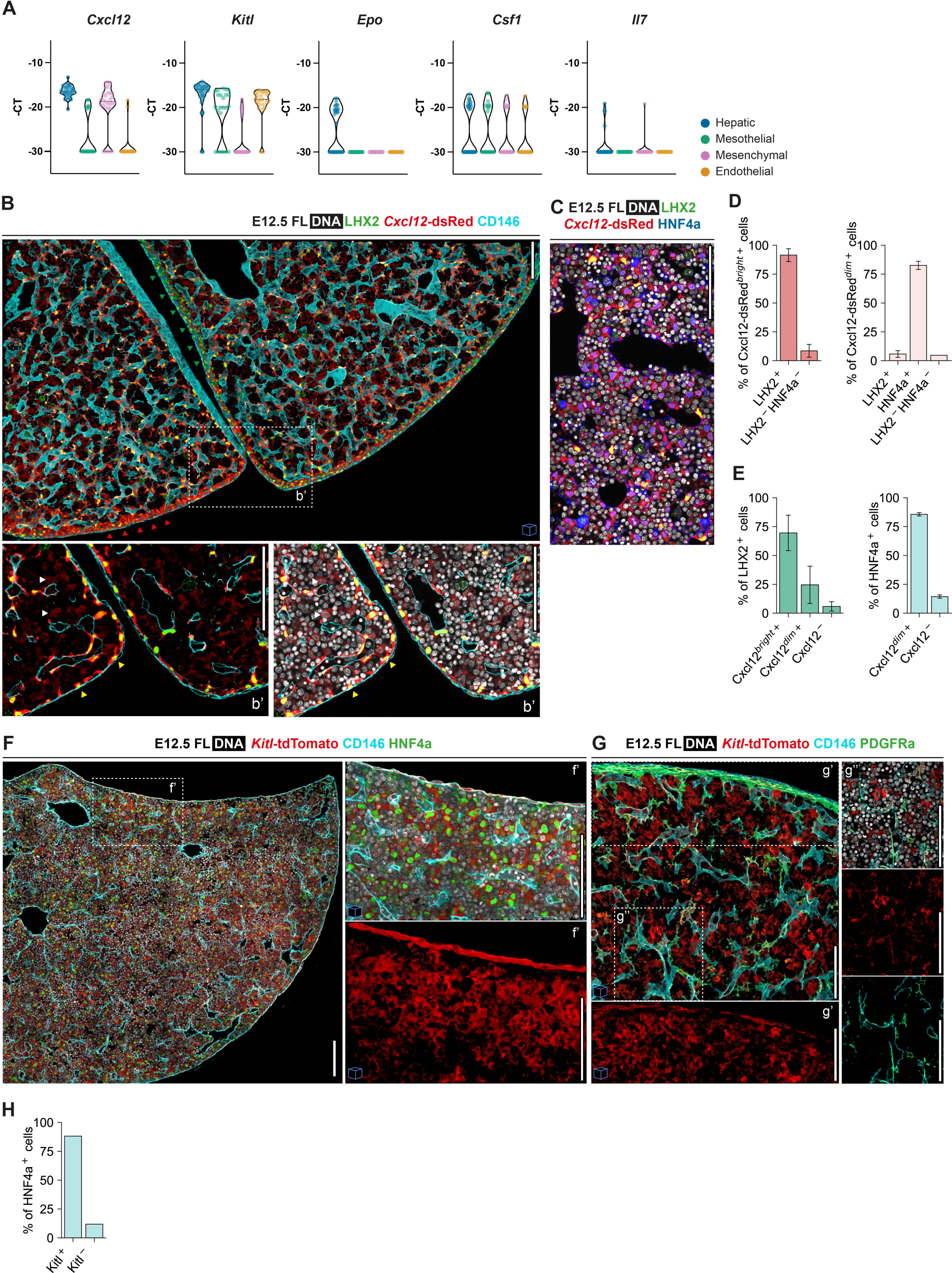
Supportive hematopoietic cytokines are produced by distinct stromal populations and accumulate at the mesothelial and sub-mesothelial regions. **(A)** Violin plots of single-cell gene expression analysis of the cytokines *Cxcl12, Kitl, Epo, Csf1* and *Il7* in Hepatic (blue), Mesothelial (green), Mesenchymal (pink) and Endothelial (orange) cells at E12.5. **(B)** IHC of E12.5 FL (*Cxcl12*-dsRed mice) with LHX2 (green), *Cxcl12*-dsRed (red) and CD146 (cyan), 3D view (25 μm). Red arrowheads point to a border region with bright *Cxcl12*-dsRed cells, while green arrowheads indicate a region with dim *Cxcl12*-dsRed cells. Insert b’ shows an enlarged single-stack view of the selected region. Yellow arrowheads indicate cells with high expression of *Cxcl12*, that co-stain LHX2. White arrowheads indicate cells with low expression of *Cxcl12*, LHX2^−^. Scale bar 100 μm. **(C)** Single-stack IHC of E12.5 FL (*Cxcl12*-dsRed mice) with LHX2 (green), *Cxcl12*-dsRed (red) and HNF4a (blue). Scale bar 100 μm. **(D)** Quantification of *Cxcl12*-dsRed^bright+^ and *Cxcl12*-dsRed^dim+^ cells in the stromal compartment. Bar plots indicate average ± standard deviation (n=6). **(E)** Quantification of Cxcl12-dsRed cells with distinct signal intensities in LHX2^+^ (left) and HNF4a^+^ (right) stromal cells. Bar plots indicate average ± standard deviation (n=6). **(F)** Single-stack IHC of E12.5 FL (*Kitl*-tdTomato mice) with HNF4a (green), *Kitl*-tdTomato (red) and CD146 (cyan). Inserts f’ show an enlarged 3D view (20 μm) of the selected regions. Scale bar 100 μm. **(G)** IHC of E12.5 FL (*Kitl*-tdTomato mice) with PDGFRa (green), *Kitl*-tdTomato (red) and CD146 (cyan), 3D view (15 μm). Inserts g’ and g’’ show an enlarged 3D view or single-stack of the selected regions, respectively. Scale bar 100 μm. **(H)** Quantification of *Kitl-t*dTomato^+^ cells in HNF4a^+^ cells. HNF4a^+^ classified cells from a single-image (3D image, 16.000 total cells) were selected and Kitl-dTomato^+^ cells manually classified (n=1).

To characterize the tissue distribution of the two most expressed cytokines, CXCL12 and KITL, we analyzed FL from reporter *Cxcl12*-dsRed^(34)^ and *Kitl*-dTomato mice^(35)^. The cells with high levels of *Cxcl12*-dsRed co-expressed LHX2 and were predominantly located in the sub-mesothelial and perivascular regions (Fig4B, D). The expression of *Cxcl12* in the sub-mesothelial region was heterogeneous, with some regions with the accumulation of cells expressing bright *Cxcl12*-dsRed (indicated by red arrowheads in Fig4B). Cells with dull expression of *Cxcl12* were found in the parenchyma, and co-expressed HNF4a (Fig4C, D). Nearly all mesenchymal cells and hepatoblasts expressed *Cxcl12* (Fig4E).

A heterogeneous pattern of *Kitl*-tdTomato expression was found in CD146^+^ mesothelial cells (Fig4A, F-f’, G-g’). *Kitl* expression was also found in HNF4a+ hepatoblasts (Fig4F) but not in endothelial and mesenchymal cells (Fig4G, H). These results indicate that distinct stromal populations can express the same cytokines expression is shared distinct stromal populations regardless of cell distribution within the tissue.

### At later stages, IE-derived but not YS-derived progenitors display a homogeneous distribution throughout the parenchyma

Given that the FL is a major site of hematopoiesis during development, we investigated the distribution of hematopoietic cells at subsequent developmental stages. We found that KIT^+^ progenitors maintained their accumulation close to the capsule (Fig5A, B) at E14,5 while CD45^+^ cells and DP progenitors changed their spatial localization. CD45^+^ cells displayed a lower frequency within 50 μm from the capsule, while DP progenitors were evenly dispersed in the parenchyma (Fig5B). Analysis of cell-cell contacts showed that as for the earlier time-points, hematopoietic progenitors (KIT+ and DP) preferentially interacted with hepatoblasts rather than with stellate cells (Fig5C).

**Figure 5.**
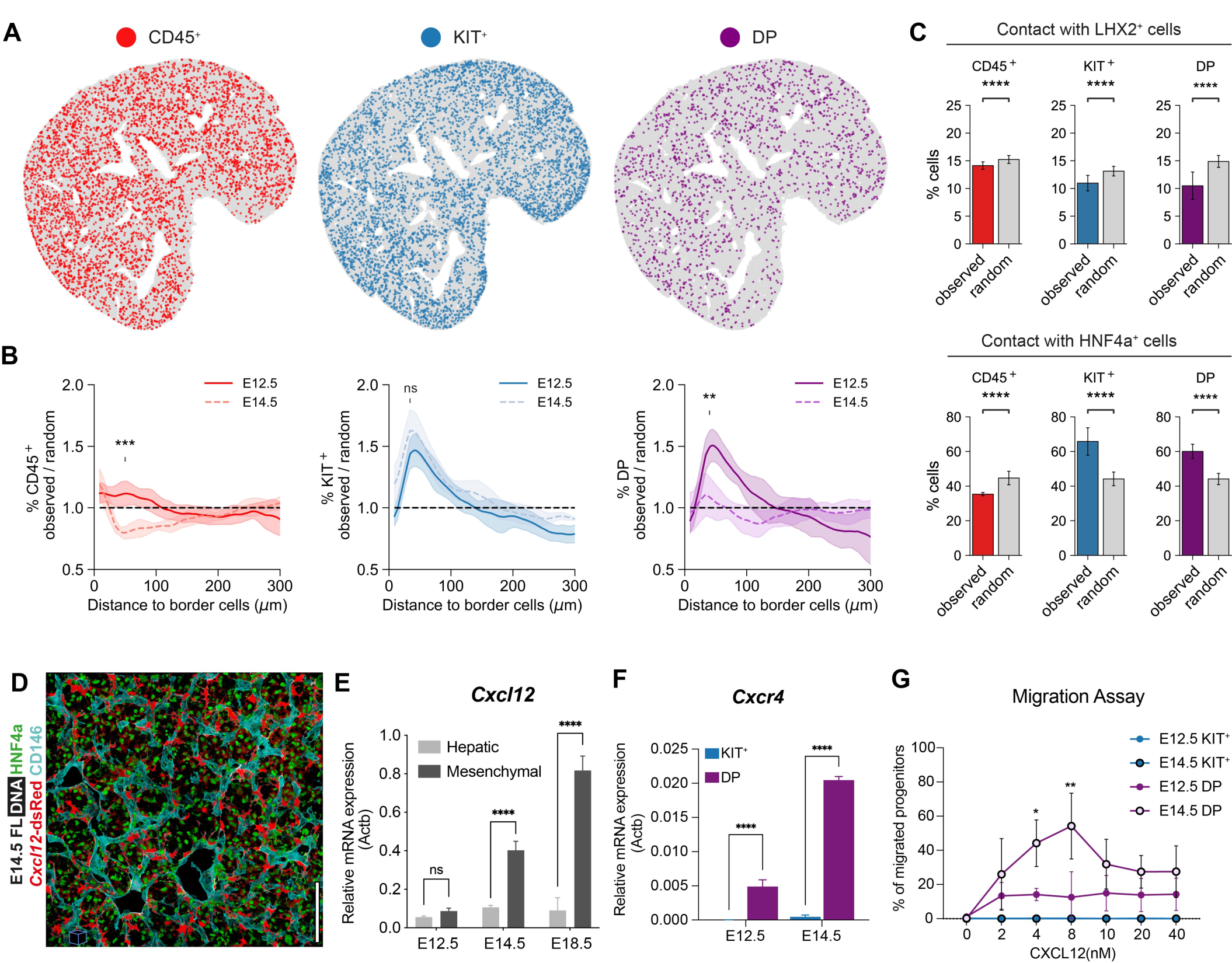
At later stages, opposite to YS-derived, IE-derived progenitors display a homogeneous distribution throughout the parenchyma in response to *Cxcl12* levels. **(A)** Dot plot representation of CD45^+^ (red), KIT^+^ (blue) and DP (purple) cell distribution of E14.5 FL representative sections (single-stack). **(B)** Distance to the border profile of CD45^+^ (red, n=6), KIT^+^ (blue, n=6) and DP (purple, n=6) cells of E14.5 FLs. E12.5 data from Figure 3D is overlaid for comparison. Curves indicate average ± standard deviation. Statistical analysis was calculated at the peak of each population using the Mann-Whitney test, between E12.5 and E14.5 curves. *, P < 0.05; **, P < 0.01; ****, P < 0.0001; ns, not significant. **(C)** Neighborhood composition of CD45^+^, KIT^+^ and DP cells of E14.5 FL. Graphs display the percentage of cells that are in contact with LHX2^+^ (left, n=3) and HNF4a^+^ (middle, n=3) stromal cells (observed) compared to random. Bar plots indicate average ± standard deviation. Statistical analysis was calculated using t-test Welch. *, P < 0.05; **, P < 0.01; ****, P < 0.0001; ns, not significant. **(D)** Single-stack IHC of E14.5 FL (*Cxcl12*-dsRed mice) with LHX2 (green), *Cxcl12*-dsRed (red) and HNF4a (blue). Scale bar 100 μm. **(E)** Quantitative RT-PCR analysis of the expression levels of the hematopoietic chemokine *Cxcl12* in hepatic (light gray), and mesenchymal (dark gray) cells in E12.5, E14.5 and E18.5 FL cells (n=3). Statistical analysis was calculated using two-way ANOVA, followed by Šídák’s multiple comparisons test. ****, P < 0.0001; ns, not significant. **(F)** Quantitative RT-PCR analysis of the expression levels of the CXCL12 receptor (*Cxcr4*) in KIT^+^ (blue), and DP (purple) cells in E12.5, E14.5 FLs (n=3). Statistical analysis was calculated using two-way ANOVA, followed by Šídák’s multiple comparisons test. ****, P < 0.0001. **(G)** Transwell migration assay of DP and KIT^+^ progenitors of E12.5 and E14.5 FL in response to the chemokine CXCL12a. Data are presented as the percentage of input progenitors that migrate to the bottom chamber (n=3). Statistical analysis was calculated using two-way ANOVA, followed by Šídák’s multiple comparisons test. *, P < 0.05; **, P < 0.01.

To investigate the mechanisms underlying the changes in spatial distribution of progenitors we analyzed the CXCL12-CXCR4 axis, involved in hematopoietic cell migration^(36)^. We observed strong *Cxcl12*-dsRed expression in perivascular cells at E14.5 (Fig5D). While hepatoblasts maintained similar *Cxcl12* transcript expression at all stages, mesenchymal cells expressed significantly higher levels with time (Fig5E). Whereas KIT^+^ progenitors have no expression of *Cxcr4*, DP progenitors express low levels of the chemokine receptor at E12.5, which were further increased at E14.5 (Fig5F). In a dual-chamber migration assay, KIT^+^ progenitors did not migrate in response to CXCL12 at either E12.5 or E14.5 whereas E12.5 DP progenitors migrate in response to CXCL12, which was more pronounced at E14.5 (Fig5G). We propose that E14.5 DP progenitors migrate from sub-capsular region into the parenchyma, between E12.5 and E14.5 in response to the redistribution of CXCL12 expressing mesenchymal cells.

At later stages of mouse development, hematopoietic cells transit from the FL to the BM, where hematopoiesis is sustained throughout adulthood. This migration to the BM starts at around E15.5-E16.5, and few progenitors are found in the FL before birth^(37,38)^. We observed an accumulation of CD45^+^ cells in the major vessels and in the vicinity of sub-mesothelial cells in E18.5 FLs (FigS5A-D). At E18.5 we found clusters of both CD11b^+^ myeloid and B220^+^ lymphoid cells around the vessels and in sub-capsular regions (FigS5E-J). CD11b^+^ cells were equally represented in major vessels and the sub-capsular region, while B220^+^ cells were primarily located at the latter (FigS5K, L). These results point to the sub-mesothelial region as a supportive site for hematopoiesis also at late developmental stages.

### The fetal liver is a low-cytokine environment

To understand the cytokine expression dynamics in the FL we compared transcript expression in all stromal populations from E12.5 up to E18.5 (FigS6A, B). Mesenchymal cells were the major producers of *Cxcl12* after E12.5, with increasing levels at later stages. Conversely, *Kitl*, *Epo* and *Il7* were predominantly expressed by hepatoblasts and exhibited a decrease in expression at E18.5, with few hepatoblasts expressing *Cxcl12* and *Kitl* at this timepoint. *Thpo*, primarily expressed by hepatoblasts, was increased at later stages. We observed higher levels of the cytokines *Flt3* and *Csf1* in mesothelial and perivascular cells at E18.5, possibly supporting the accumulation of lymphoid and myeloid cells in these regions.

We compared of the highest stromal producer in the FL with BM endothelial and CD51^+^CD106^+^ cells (a mesenchymal stromal population reported to be highly enriched for hematopoietic cytokines)^(39)^, from postnatal days 7, 14, and 21 to better understand the differences in cytokine expression (Fig6A; FigS6C). Except for *Epo*, known to be produced in the post-natal kidney, all cytokines were expressed at significantly higher levels in the BM. Moreover, we observed that CD51^+^CD106^+^ cells were the primary cytokine producers, in neonatal BM (Fig6A) where *Cxcl12*, *Kitl*, and *Il7* transcripts were more than 10 times higher than in FL. These observations indicate that the FL is a low-cytokine environment.

## Discussion

We analyzed FL hematopoiesis between E12.5 and E14.5 because it corresponds to the period when HSC are found in increasing numbers^(20)^. Although the degree of expansion of HSC in FL is still under debate^(40,41)^, understanding how the FL stromal compartment regulates fetal hematopoiesis is essential to understand the mechanism leading to HSC formation and to the pathogenesis of childhood hematologic disorders.

The comparative analysis of all non-hematopoietic stromal subsets at E12.5 showed the hepatoblasts as major producers of *Kitl* and *Cxcl12* and the only cells that expressed *Epo* and *Il7*. Cytokine production decreases by E18.5 consistent with previous reports^(29)^. *Kitl*, however, is also produced by mesothelial and endothelial cells, whereas *Cxcl12* is found in stellate cells/pericytes and NG2 peri-portal cells. This promiscuous distribution of two major hematopoietic modulators explains why inhibiting cytokine production in defined cell types had limited impact in hematopoiesis and in the HSC FL compartment. The NG2 expressing cells are found, in FL, only around large portal vessels rather than around sinusoids and they have been proposed to play important roles in fetal hematopoiesis^(25)^. However, NG2 expressing cells are rarely found at the early stages of active hematopoiesis (E12.5) and later (E14.5) they accumulate around the nascent portal network where cholangiocytes differentiate. NG2 is also expressed in mesothelial cells that we identify as a new element of the FL stroma, constituting the FL capsule and producing *Kitl*. Disrupting NG2 expression might, therefore, have unpredictable consequences for the integrity of the mesothelial compartment and for the organ architecture and could explain the modest decrease in HSC, previously reported^(25)^. Proximity to NG2+ cells was previously evaluated for 105 cells from 12 E14.5 FL (an average of 9 cells per FL that contains more than 1000 HSC), a number that may not be representative of the entire HSC compartment at that stage. Constitutive deletion of KITL in both endothelial and PDGFRa expressing cells significantly impacted HSC numbers but only postnatally^(29)^, indicating that KITL produced by these two cell subsets did not impact the HSC compartment before birth. DLK1 expression has been considered a marker for hepatoblasts, however, we found it expressed in lymphatic endothelial cells, mesothelial and stellate cells.

At E12.5, the sub-mesothelial thick layer of stellate cells producie *Cxcl12*, but not *Kitl.* Proliferating hepatoblasts also concentrate in this area where they receive signals from mesothelial cells to expand (hepatocyte growth factor, midkine and pleiotropin)^(42)^. Mesothelial cells and hepatoblasts are the main producers of KITL and hematopoietic progenitors of YS and IE origin converge to this region, where they are in direct contact with hepatoblasts, but not with endothelial or stellate cells. At this stage, this FL region is also highly hypoxic which might contribute to the highest levels of Epo detected in hepatoblasts^(43)^. At E14.5, CXCL12-producing stellate cells have invaded the parenchyma possibly recruited by the expansion of the vascular network, and NG2^+^ peri-vascular cells accumulate around the portal vessels. The sub-mesothelial layer of stellate cells is reduced and IE hematopoietic progenitors no longer concentrate in this region. At this stage, *Cxcl12* expression by stellate cells increased and the levels of *Cxcr4* in hematopoietic progenitors also increased with augmented response to chemokine gradients. YS-derived progenitors that comprise mostly erythroid cells do not express *Cxcr4* and are still found in the sub-mesothelial region, at E14.5. Later, at E18.5, hematopoietic cytokines are no longer produced by hepatoblasts, that become rare. Cholangiocytes accumulate around portal vessels close to NG2 expressing cells and where CD45^+^ B220^+^ or CD11b^+^ cells now converge. B220^+^ and CD11b^+^ lymphoid and myeloid cells respectively also accumulate in the sub-capsular region. Stellate cells that also surround sinusoids are likely to drive the exit of mature cells and progenitors from FL, after E15.5.

The modest impact observed on fetal hematopoiesis by deleting cytokines or chemokines in specific cell types can be due to the redundant production of these factors, as discussed above. It could, however, also indicate that HSC are largely independent from these factors before BM colonization. We found that YS-derived erythroid progenitors outcompete HSC derived progenitors, for access to low concentrations of EPO present in FL, because they have a lower threshold response to this factor^(5)^. Moreover, the low concentration of IL-7 in the FL allows B1 B cell selection for autoreactive specificities^(44)^. We show here that the FL is generally a low cytokine environment compared to BM. It is therefore possible that YS or IE-derived cells endowed with low HSC potential, previously reported^(45,46)^, are particularly efficient in consuming the limited environmental resources available. HSC or their progenitors could complete a limited number of rounds of division in the FL, in a cytokine independent manner after migrating from in the dorsal aorta. Their higher threshold for cytokine response prevents differentiation and explains their limited contribution to fetal hematopoiesis^(47)^. In the present study we did not analyze HSC, so it is possible that DP cells that neighbor hepatoblasts are the progeny of multipotent cells that do not integrate the stem cell compartment. Because expansion in the FL might be limited, HSC expansion would occur preferentially in the BM shortly after colonization, as suggested^(41)^. In this scenario the FL HSC niche properties would have to be revised: it would be a cytokine protected environment rather than being a cytokine rich environment.

## Acknowledgements

We thank A. Bandeira, P. Vieira, E. Maikranz, and P. Pereira and the members of the P. Pinto-do-Ó, C. Baroud and A. Cumano laboratories for critical discussions. We thank C. Ait-Mansour (SONY Europe B.V) for help with spectral cytometry and critical discussions. We thank D. Hardy (Institut Pasteur, Paris, France) for the FLK1-GFP reporter mice. We acknowledge the Center for Translational Science (CRT) - Cytometry and Biomarkers Unit of Technology and Service (CB UTechS) at Institut Pasteur for support in conducting this study, namely S. Novault, S. Megharba, S. Schmutz, V. Libri, and T. Stephen; and the staff of the animal facility of the Institut Pasteur for mouse care. We acknowledge the Histology Platform, Image Analysis Hub, The Cluster Team and Gpulab of Institut Pasteur for technical support. We gratefully acknowledge the UtechS Photonic BioImaging (Imagopole), C2RT, Institut Pasteur, supported by the French National Research Agency (France BioImaging; ANR-10–INSB–04; Investments for the Future). We acknowledge Investissement d’Avenir grant ANR-16-CONV-0005 for funding computing resources used for simulations. We acknowledge the support of i3S facilities: Animal Facility, Translational Cytometry Unit, HEMS (Histology and Electron Microscopy), and ALM (Advanced Light Microscopy), the latter, member of the national infrastructure PPBI-Portuguese Platform of BioImaging (supported by POCI-01-0145-FEDER-022122).

## Author contributions

Conceptualization: F.S.S., M.M.P., P.P.O., A.C.; Methodology: F.S.S., M.M.P., V.B., G.R.; Software: M.M.P., V.B., G.R.; Resources: X.F, J.P.P., E.A., G.A., M.d.B.; Formal Analysis: F.S.S., M.M.P., V.B.; Investigation: F.S.S., M.M.P., R.F.S., M.P.M.; Writing - Original Draft: F.S.S., A.C.; Writing - Review and Editing: F.S.S., M.M.P., V.B., A.C., P.P.O., C.B.; Visualization: F.S.S., M.M.P., V.B.; Supervision: A.C., P.P.O., C.N.B.; Funding Acquisition: A.C., P.P.O., C.N.B..

## Data availability

All raw data is available in the open repository Zenodo with the following unique identifiers: microscopy images-doi: 10.5281/zenodo.7866578; gene Expression-doi: 10.5281/zenodo.7867188; flow cytometry-doi: 10.5281/zenodo.7867212.

## Supplementary Figure legends

**Figure S1 (related to Figure 1).**
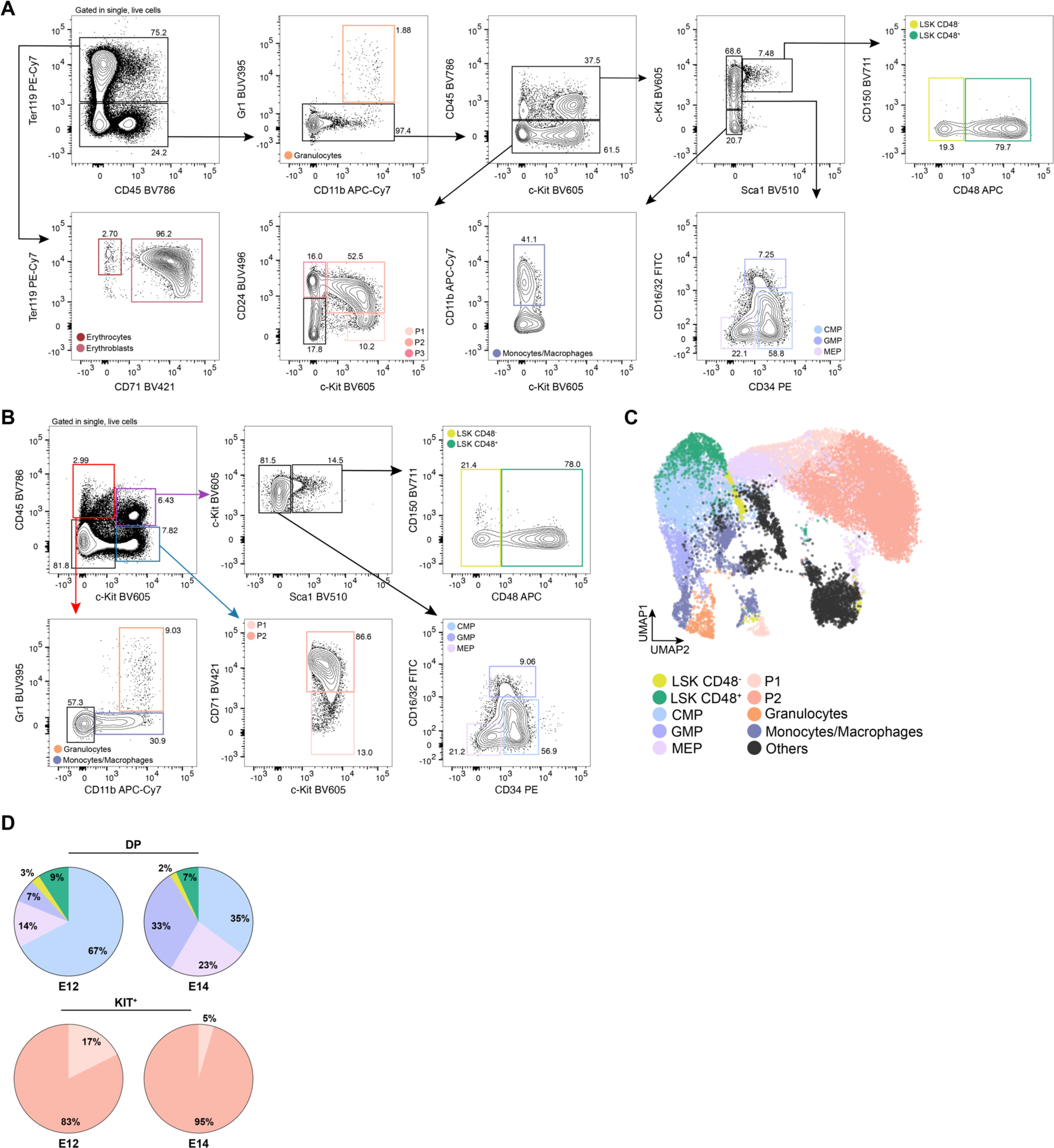
Discrimination of the hematopoietic compartment. **(A)** Gating strategy used for the analysis of the hematopoietic compartment of E12.5 FLs using the surface markers Ter119, CD45, CD71, Gr1, CD11b, KIT, Sca1, CD16/32, CD34, CD150 and CD48. (Related to Figure 1B). **(B)** Gating strategy and **(C)** UMAP analysis showing the cell types that are included in the image analysis with CD45 and KIT. **(C)** Percentage of the progenitor cell types within DP cells (top) and KIT^+^ cells (bottom) at E12 (left) or E14 (right).

**Figure S2 (related to Figure 1).**
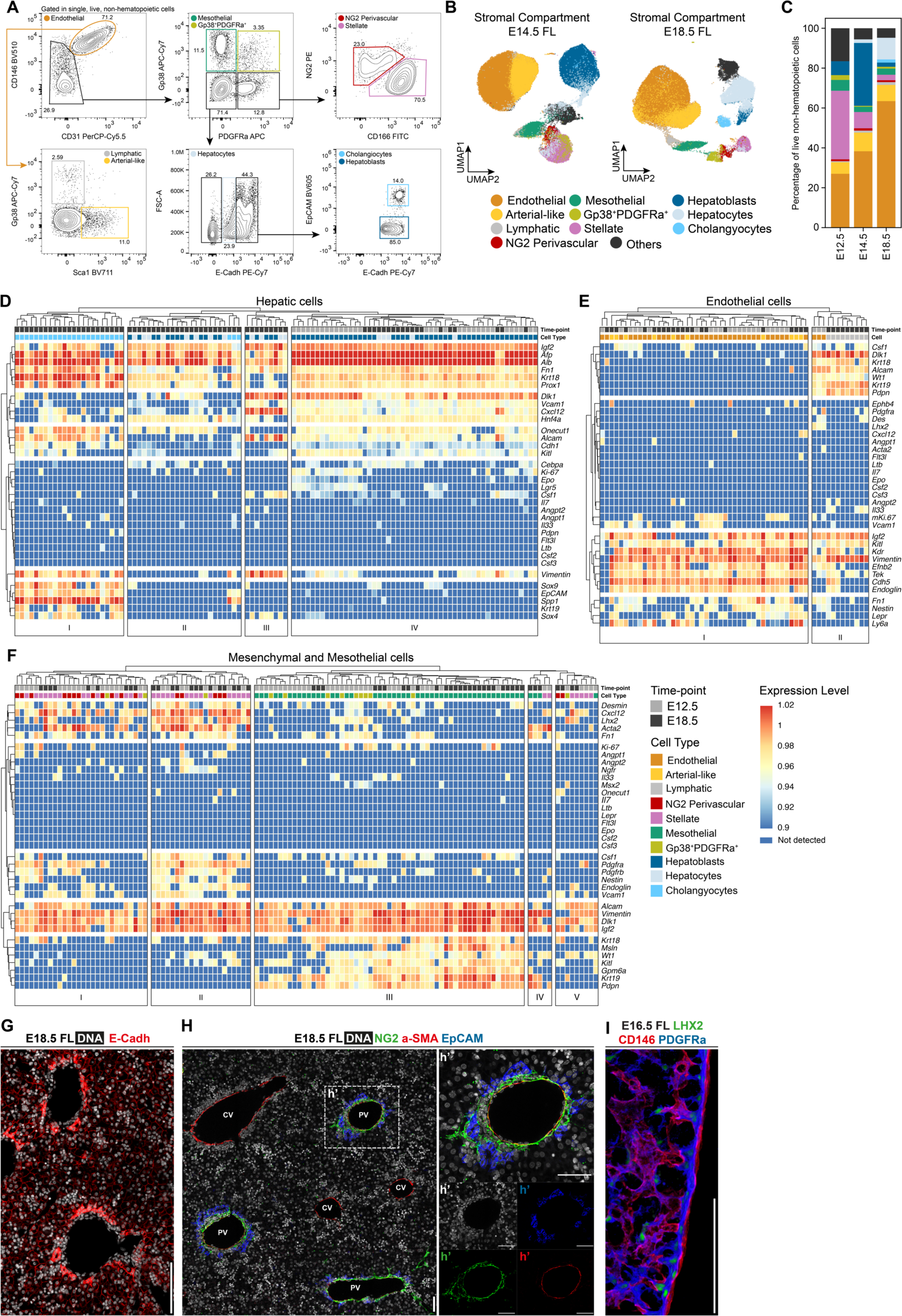
Characterization of the stromal compartment. **(A)** Gating strategy used for the analysis of of Ter119^−^CD45^−^CD71^−^KIT^−^ FL cells using the surface markers CD146, CD31, Gp3, PDGFRa, NG2, CD166, Sca1, E-Cadh and EpCAM. Represented is the profile of an E18.5. The same strategy was applied to all time-points. **(B)** UMAP analysis of flow cytometry data of Ter119^−^CD45^−^CD71^−^KIT^−^ cells from E14.5 (left) and E18.5 (right) FLs stained with the surface markers CD146, CD31, Gp3, PDGFRa, NG2, CD166, Sca1, E-Cadh and EpCAM. **(C)** Distribution profile of live non-hematopoietic cells at E12.5, E14.5 and E18.5 as obtained from flow cytometry data. **(D)** Heatmap of single-cell multiplex qPCR in sorted hepatic cells of E12.5 and E18.5 FLs. Each column represents a single cell, and it is color-coded according to time-point (E12.5, E18.5) and cell type (Hepatoblasts, Hepatocytes and Cholangiocytes). Gene expression was normalized to *β-actin* and *Gapdh*, and unsupervised hierarchical clustering was performed. **(E)** Heatmap of single-cell multiplex qPCR in sorted endothelial cells of E12.5 and E18.5 FLs. Each column represents a single cell, and it is color-coded according to time-point (E12.5, E18.5) and cell type (Endothelial, arterial-like and lymphatic). Gene expression was normalized to *β-actin* and *Gapdh*, and unsupervised hierarchical clustering was performed. **(F)** Heatmap of single-cell multiplex qPCR in sorted mesenchymal and mesothelial cells of E12.5 and E18.5 FLs. Each column represents a single cell, and it is color-coded according to time-point (E12.5, E18.5) and cell type (Mesothelial, Gp38^+^PDGFRa^+^, NG2 Perivascular and Stellate). Gene expression was normalized to *β-actin* and *Gapdh*, and unsupervised hierarchical clustering was performed. **(G)** Single-stack IHC of E18.5 FL with DAPI (white) and E-Cadherin (red). Scale bar 100 μm. **(H)** Single-stack IHC of E18.5 FL with DAPI (white), NG2 (green), aSMA (red) and EpCAM (blue). Insert h’ shows an enlarged view of a portal vessel and the respective single-channels below. Scale bar 50 μm. **(I)** 3D view (25 μm) of IHC of E16.5 with LHX2 (green), CD146 (red), and PDGFRa (blue). Scale bar 50 μm.

**Figure S3 (related to Figure 2).**
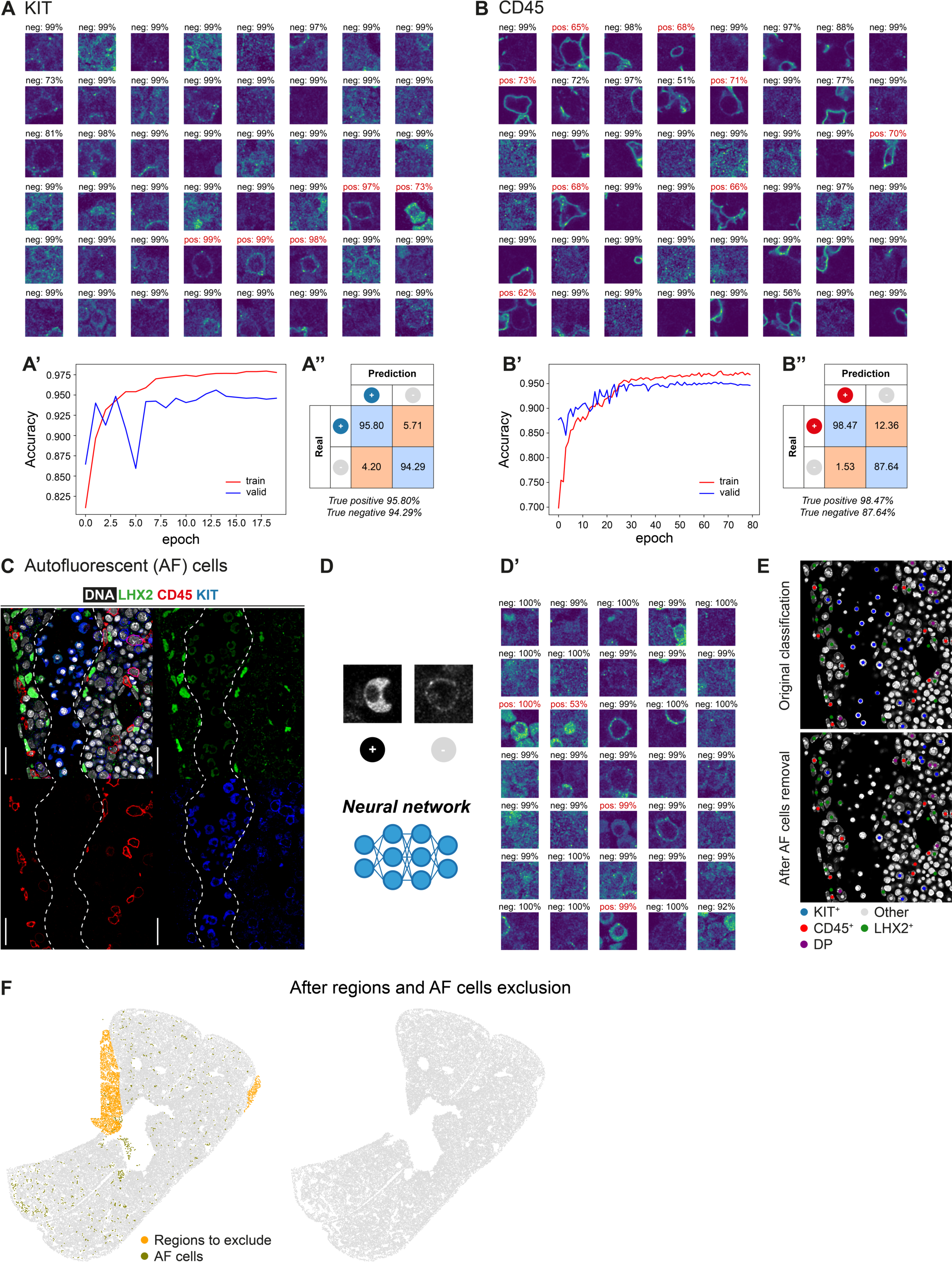
Validation and quality control of image analysis. **(A)** Validation of KIT classification. Representative images of single cells classification and respective classification confidence. (A’) Accuracy of the classification during network training on training and validation datasets. (A’’) Confusion matrix on the validation dataset. **(B)** Validation of CD45 classification. Representative images of single cells post-classification and respective classification confidence. (B’) Accuracy of the classification during network training on training and validation datasets. (B’’) Confusion matrix on the validation dataset. **(C)** Single-stack IHC of E12.5 FL with DAPI (white), LHX2 (green), CD45 (red), and KIT (blue), to highlight a region where nucleated erythrocytes (autofluorescent - AF - cells) can be seen inside a blood vessel (represented by the dashed line). **(D)** AF cells are manually classified as positive and negative to train a neural network, so that negative cells can be later removed from the analysis. (D’) Representative images of single cells classification and respective classification confidence. **(E)** DAPI channel of the tile depicted in **(C)** overlaid with classification before (top) and after (bottom) removing AF cells. **(F)** FL section (same as in Figure 2) before and after removing regions manually selected using *Coloriage*, and AF cells, highlighted on the left panel at orange and olive green, respectively.

**Figure S4 (related to Figure 3).**
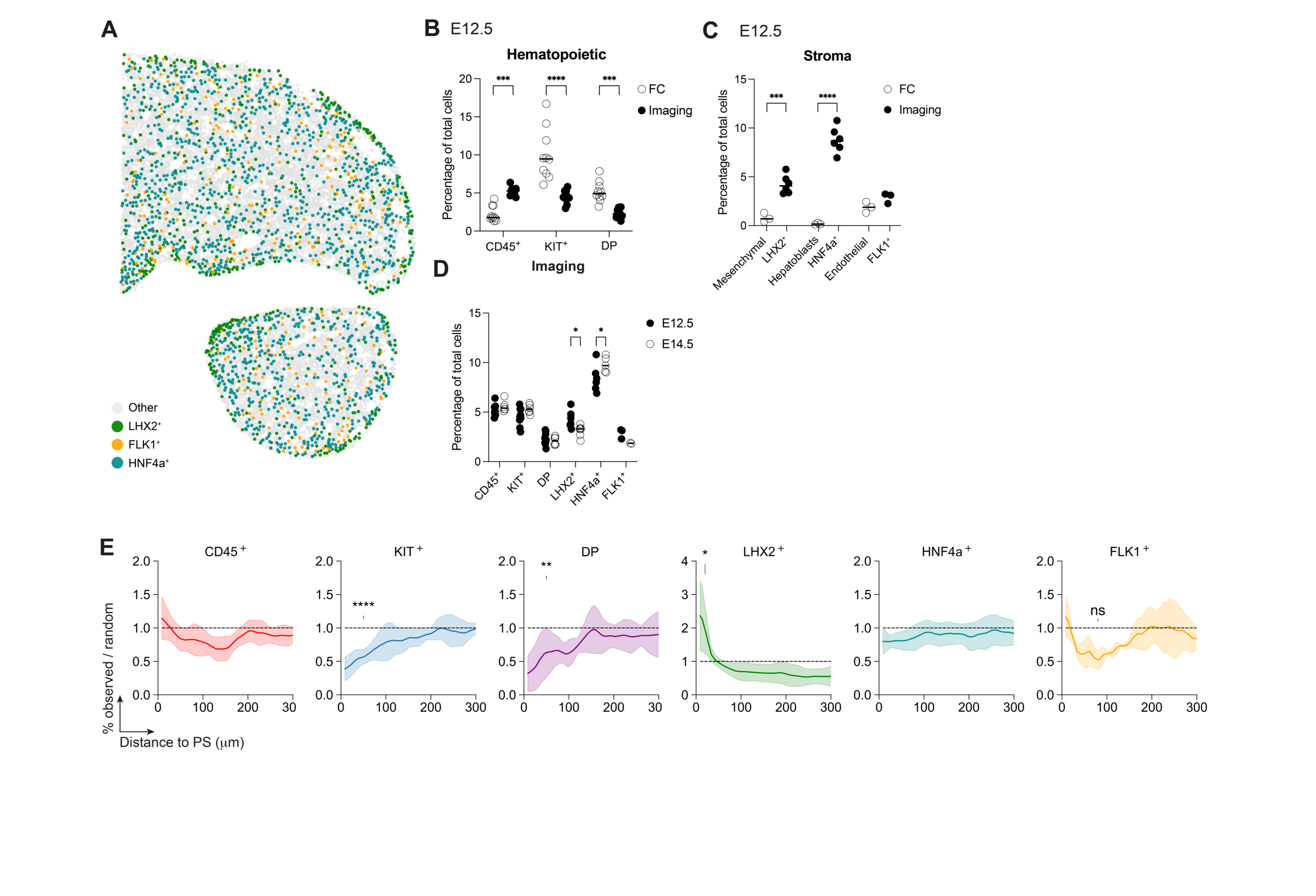
**(A)** Dot plot representation of LHX2^+^ (green), HNF4a^+^ (cyan) and FLK1^+^ (orange) cells distribution of E12.5 FL section represented in Figure 3 (10 μm thick projected into a single plane). **(B)** Percentage of hematopoietic cells recovered and analyzed by flow cytometry (n=9) versus quantified by imaging of FL sections (n=9). Statistical analysis was calculated using two-way ANOVA, followed by Šídák’s multiple comparisons test. ***, P < 0.001; ****, P < 0.0001. **(C)** Percentage of stromal cells recovered and analyzed by flow cytometry (n=3) versus quantified by imaging of FL sections (LHX2 and HNF4a (n=6), FLK1 (n=3)). Statistical analysis was calculated using two-way ANOVA, followed by Šídák’s multiple comparisons test. ***, P < 0.001; ****, P < 0.0001. **(D)** Percentage of hematopoietic and stromal cells quantified by imaging of FL sections at E12.5 and E14.5 (<u>E12.5</u>: CD45, KIT, DP (n=10); LHX2, HNF4a (n=6); FLK1 (n=3); <u>E14.5</u>: CD45, KIT, DP (n=6); HNF4a (n=5); LHX2 (n=6); FLK1 (n=2)). Statistical analysis was calculated using two-way ANOVA, followed by Šídák’s multiple comparisons test. *, P < 0.05. **(E)** Distance to *portal sinus* (PS) profile of CD45+ (red, n=10), KIT^+^ (blue, n=10), DP (purple, n=10), LHX2+ (green, n=6), HNF4a+ (cyan, n=6) and FLK1+ (orange, n=3) cells of E12.5 FL sections. Curves indicate average ± standard deviation. Statistical analysis was calculated at the peak of each population using the Mann-Whitney test, between observed and random curves. *, P < 0.05; **, P < 0.01; ****, P < 0.0001; ns, not significant.

**Figure S5.**
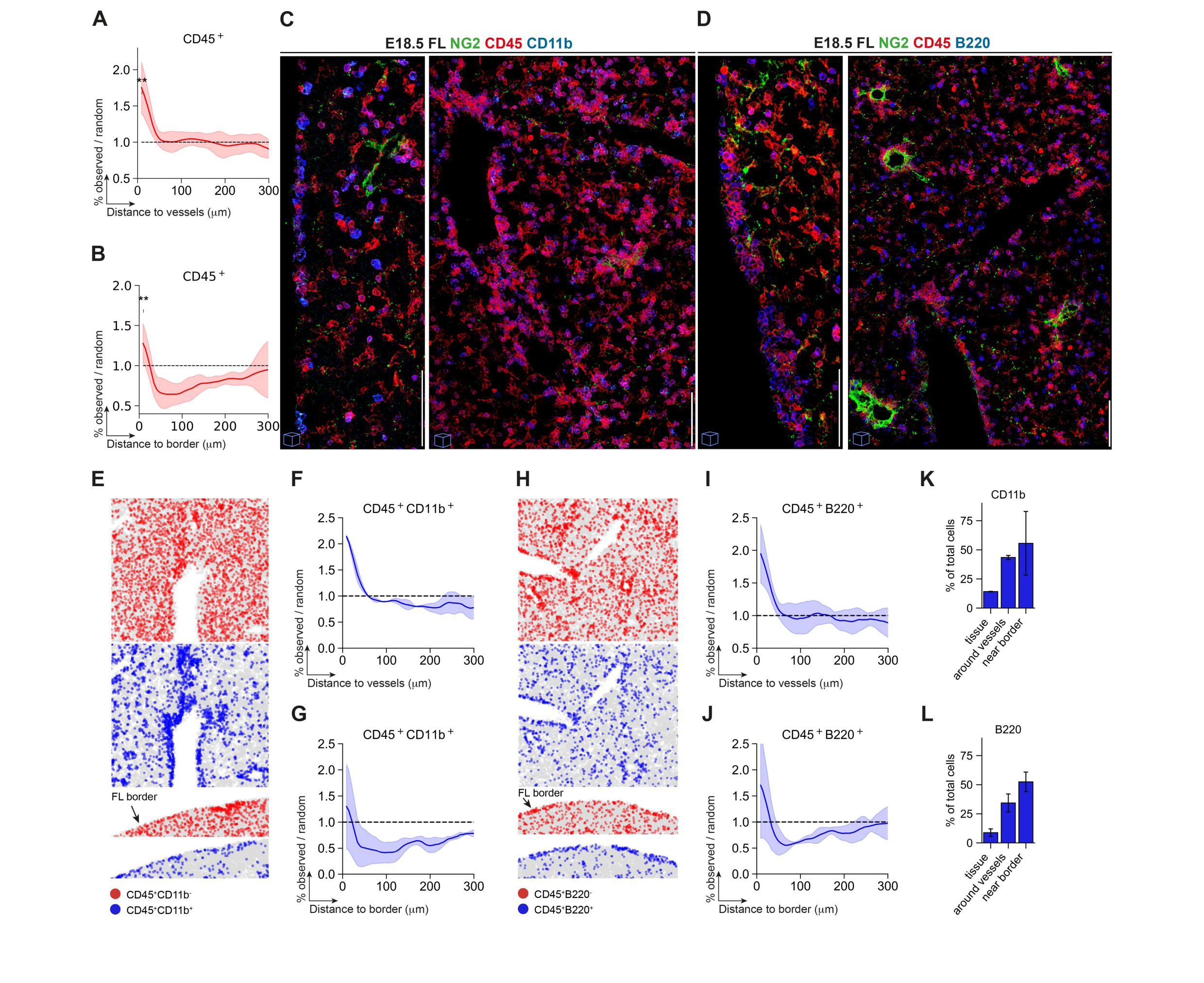
Myeloid and lymphoid cells accumulate in the vessels and mesothelium at late gestation. **(A)** Distance to vessels profile of CD45^+^ cells of E18.5 FLs. Curves indicate average ± standard deviation (n=8). **(B)** Distance to the border profile of CD45^+^ cells of E18.5 FLs. Curves indicate average ± standard deviation (n=6). **(C)** 3D view (20 μm) of IHC of E18.5 mesothelial region (left) or parenchyma (right) with NG2 (green), CD45 (red) and CD11b (blue). Scale bar 50 μm. **(D)** Distance to vessels profile of CD45^+^CD11b^+^ cells of E18.5 FLs. Curves indicate average ± standard deviation (n=2). **(E)** Example of a dot plot representation of CD45^+^CD11b^−^ (red) and CD45^+^CD11b^+^ (blue) cells around vessels (top) and in the FL border (bottom) at E18.5. **(F)** Distance to vessels profile of CD45^+^CD11b^+^ cells of E18.5 FLs. Curves indicate average ± standard deviation (n=2). **(G)** Distance to the border profile of CD45^+^CD11b^+^ cells of E18.5 FLs. Curves indicate average ± standard deviation (n=2). **(H)** Example of a dot plot representation of CD45^+^B220^−^ (red) and CD45^+^B220^+^ (blue) cells around vessels (top) and in the FL border (bottom) at E18.5. **(I)** Distance to the vessels profile of CD45^+^B220^+^ cells of E18.5 FLs. Curves indicate average ± standard deviation (n=4). **(J)** Distance to the border profile of CD45^+^B220^+^ cells of E18.5 FLs. Curves indicate average ± standard deviation (n=3). **(K)** CD45^+^CD11b^+^ frequency of total cells considering the entire sections analyzed (“tissue”), the regions around the vessels (‘around vessels’) or near the border (“near border”). **(L)** CD45^+^B220^+^ frequency of total cells considering the entire sections analyzed (“tissue”), the regions around the vessels (‘around vessels’) or near the border (“near border”). Regions “around the vessels” or “near border” (3-4 cell layers) were manually selected using *Coloriage*.

**Figure S6 (related to Figure 6).**
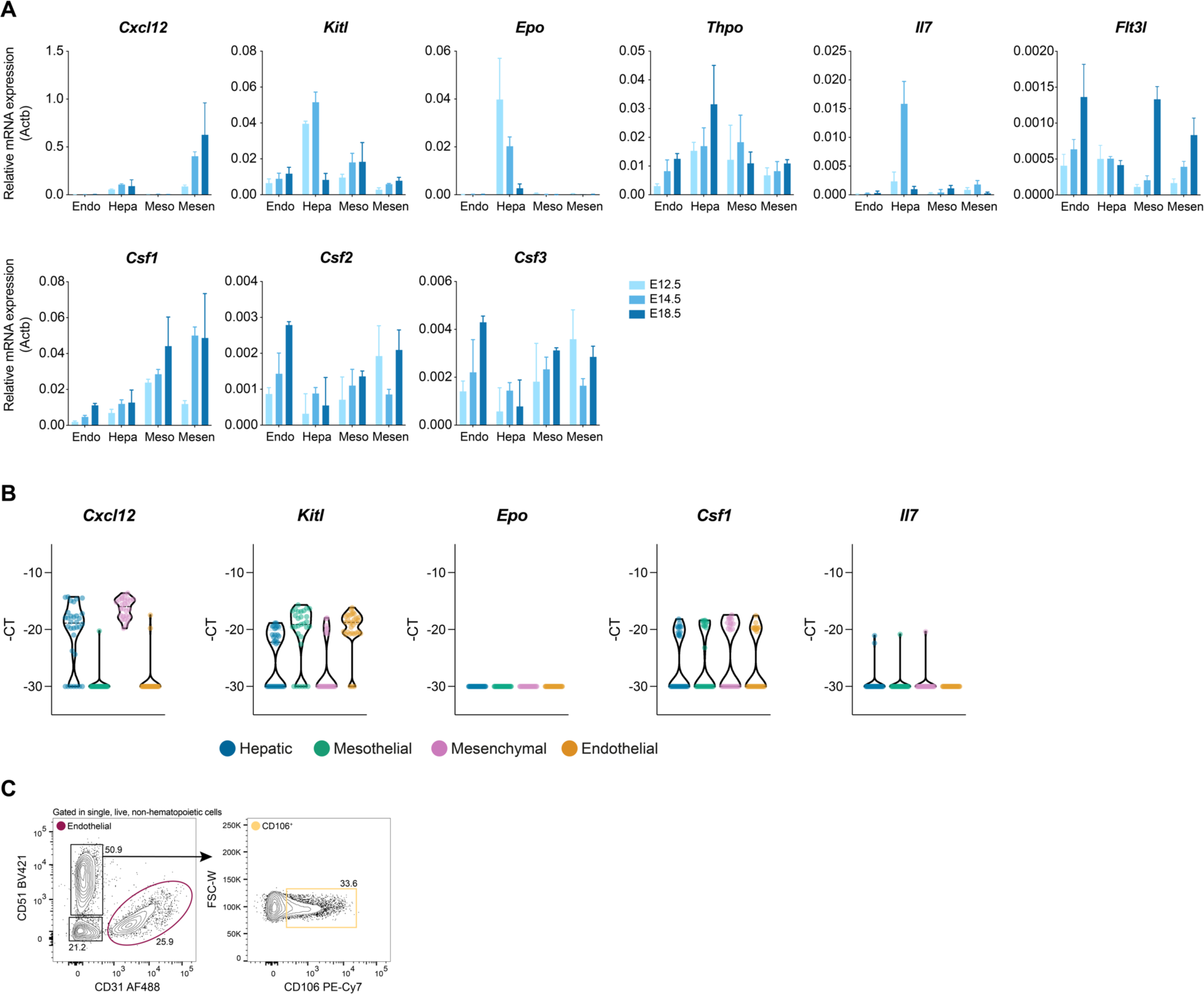
Cytokines expression along time. **(A)** Quantitative RT-PCR analysis of the expression levels of the hematopoietic cytokines *Cxcl12, Kitl, Epo, Thpo, Il7, Flt3l, Csf1, Csf2* and *Csf3* in endothelial (Endo), hepatic (Hepa), mesothelial (Meso), and mesenchymal (Mesen) cells in E12.5 (light blue), E14.5 (blue) and E18.5 (dark blue) FL cells (n=3). **(B)** Violin plots of single-cell gene expression analysis of the cytokines *Cxcl12, Kitl, Epo, Csf1* and *Il7* in Hepatic (blue), Mesothelial (green), Mesenchymal (pink) and Endothelial (orange) cells at E18.5. **(C)** Gating strategy used for the analysis of the stroma of P7, P14 and P21 BM using the surface markers CD51, CD31 and CD106. Representative profile of a P7 BM. The same strategy was applied to all time-points.

**Table S1.**
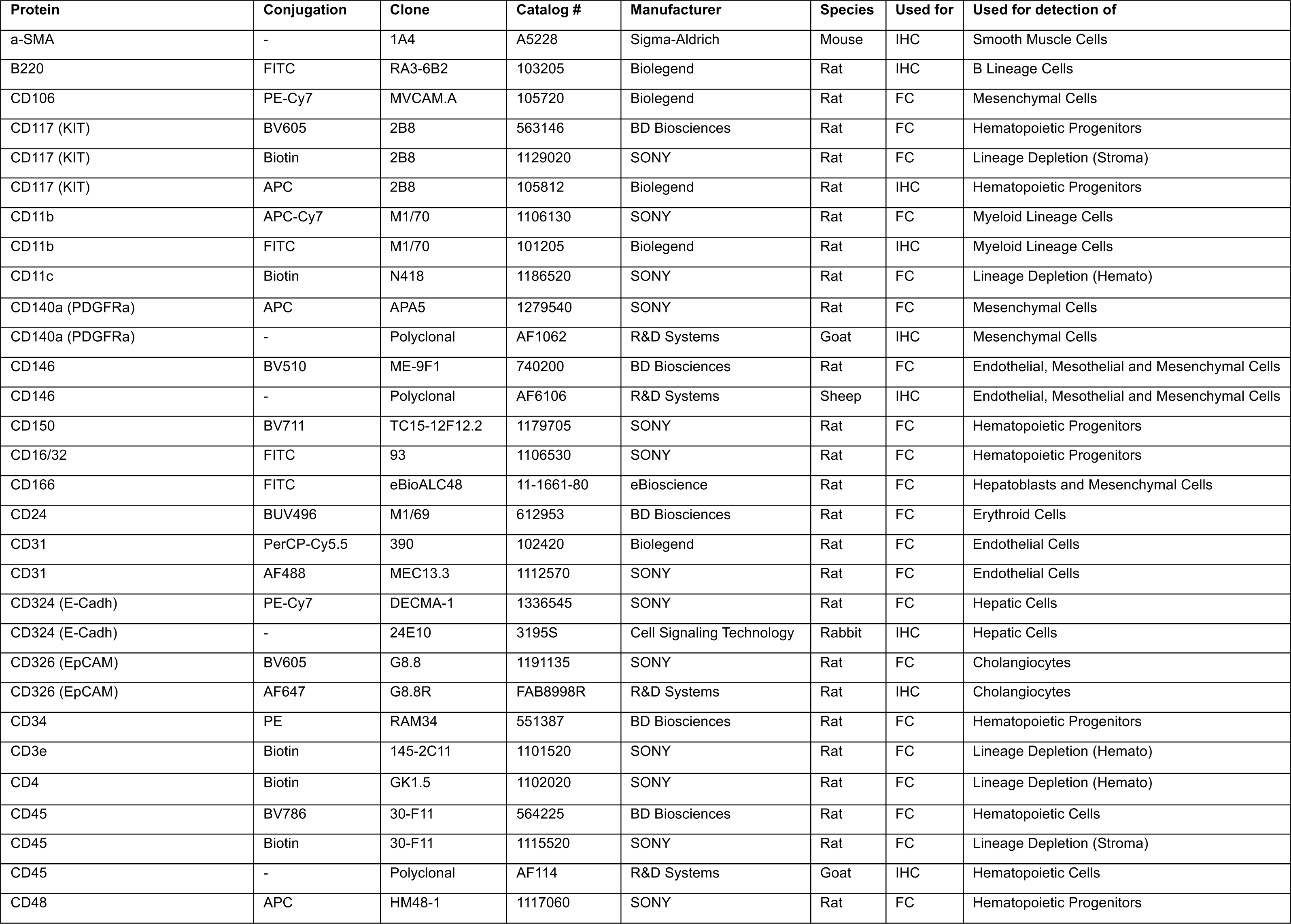
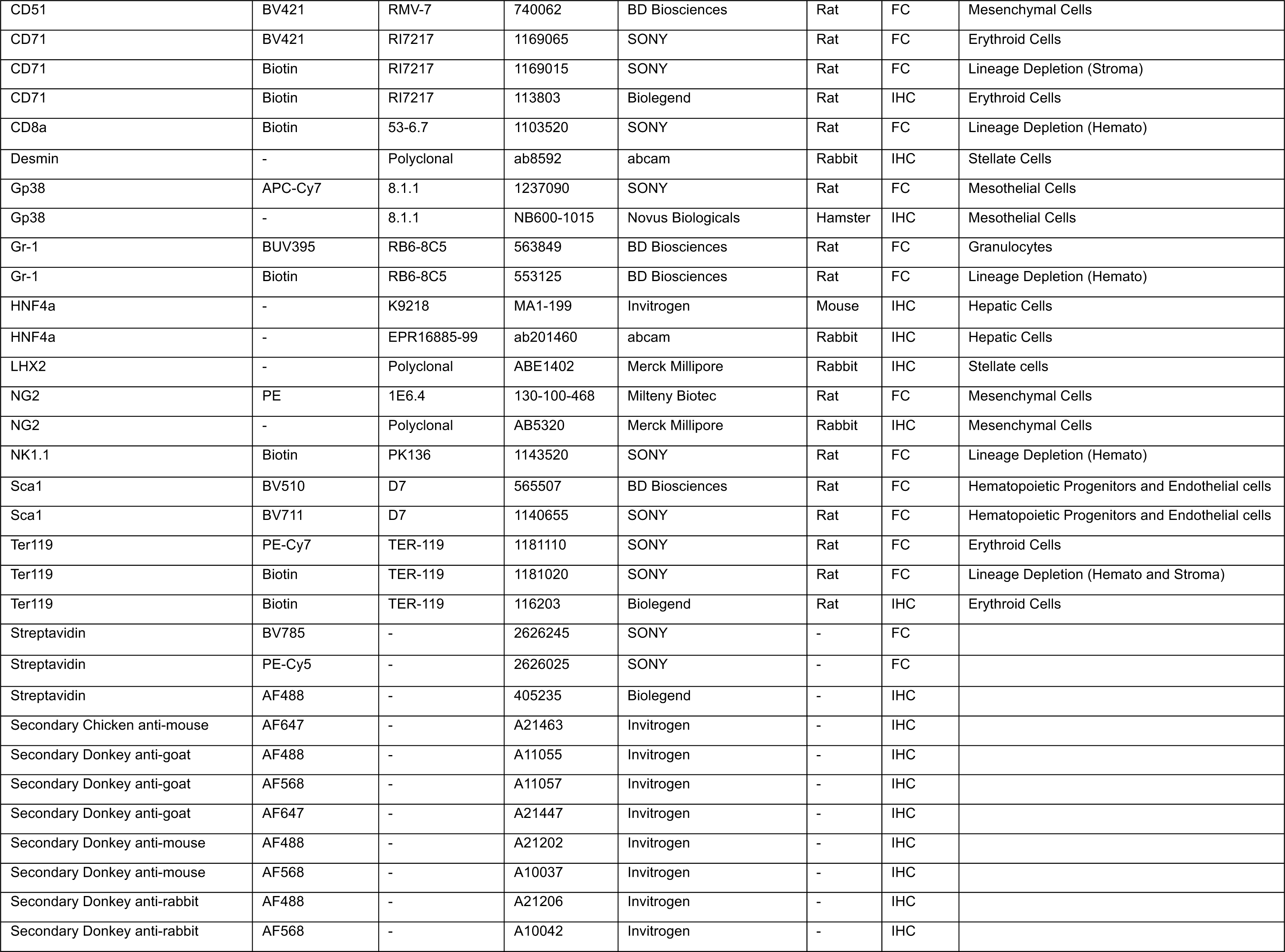
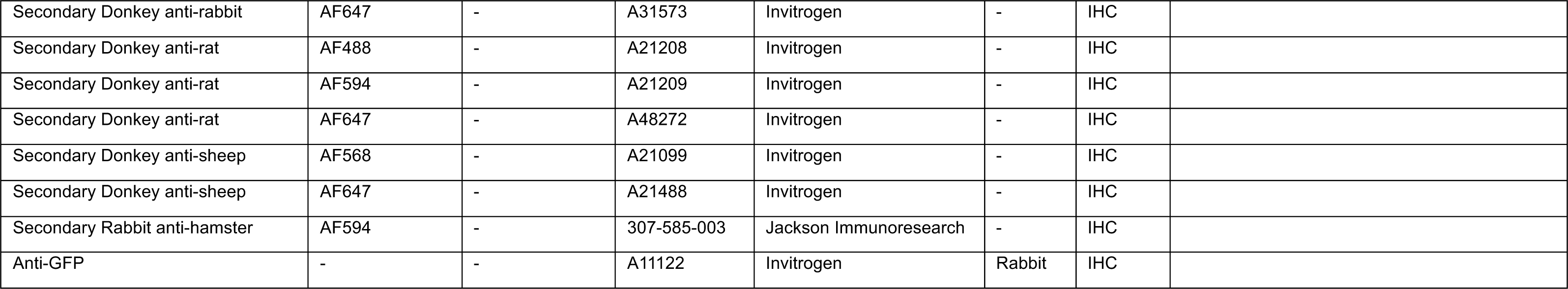
– List of antibodies for Flow Cytometry (FC) and Immunostaining (IHC)

| Protein | Conjugation | Clone | Catalog # | Manufacturer | Species | Used for | Used for detection of |
| --- | --- | --- | --- | --- | --- | --- | --- |
| a-SMA | - | 1A4 | A5228 | Sigma-Aldrich | Mouse | IHC | Smooth Muscle Cells |
| B220 | FITC | RA3-6B2 | 103205 | Biolegend | Rat | IHC | B Lineage Cells |
| CD106 | PE-Cy7 | MVCAM.A | 105720 | Biolegend | Rat | FC | Mesenchymal Cells |
| CD117 (KIT) | BV605 | 2B8 | 563146 | BD Biosciences | Rat | FC | Hematopoietic Progenitors |
| CD117 (KIT) | Biotin | 2B8 | 1129020 | SONY | Rat | FC | Lineage Depletion (Stroma) |
| CD117 (KIT) | APC | 2B8 | 105812 | Biolegend | Rat | IHC | Hematopoietic Progenitors |
| CD11b | APC-Cy7 | M1/70 | 1106130 | SONY | Rat | FC | Myeloid Lineage Cells |
| CD11b | FITC | M1/70 | 101205 | Biolegend | Rat | IHC | Myeloid Lineage Cells |
| CD11c | Biotin | N418 | 1186520 | SONY | Rat | FC | Lineage Depletion (Hemato) |
| CD140a (PDGFRa) | APC | APA5 | 1279540 | SONY | Rat | FC | Mesenchymal Cells |
| CD140a (PDGFRa) | - | Polyclonal | AF1062 | R&D Systems | Goat | IHC | Mesenchymal Cells |
| CD146 | BV510 | ME-9F1 | 740200 | BD Biosciences | Rat | FC | Endothelial, Mesothelial and Mesenchymal Cells |
| CD146 | - | Polyclonal | AF6106 | R&D Systems | Sheep | IHC | Endothelial, Mesothelial and Mesenchymal Cells |
| CD150 | BV711 | TC15-12F12.2 | 1179705 | SONY | Rat | FC | Hematopoietic Progenitors |
| CD16/32 | FITC | 93 | 1106530 | SONY | Rat | FC | Hematopoietic Progenitors |
| CD166 | FITC | eBioALC48 | 11-1661-80 | eBioscience | Rat | FC | Hepatoblasts and Mesenchymal Cells |
| CD24 | BUV496 | M1/69 | 612953 | BD Biosciences | Rat | FC | Erythroid Cells |
| CD31 | PerCP-Cy5.5 | 390 | 102420 | Biolegend | Rat | FC | Endothelial Cells |
| CD31 | AF488 | MEC13.3 | 1112570 | SONY | Rat | FC | Endothelial Cells |
| CD324 (E-Cadh) | PE-Cy7 | DECMA-1 | 1336545 | SONY | Rat | FC | Hepatic Cells |
| CD324 (E-Cadh) | - | 24E10 | 3195S | Cell Signaling Technology | Rabbit | IHC | Hepatic Cells |
| CD326 (EpCAM) | BV605 | G8.8 | 1191135 | SONY | Rat | FC | Cholangiocytes |
| CD326 (EpCAM) | AF647 | G8.8R | FAB8998R | R&D Systems | Rat | IHC | Cholangiocytes |
| CD34 | PE | RAM34 | 551387 | BD Biosciences | Rat | FC | Hematopoietic Progenitors |
| CD3e | Biotin | 145-2C11 | 1101520 | SONY | Rat | FC | Lineage Depletion (Hemato) |
| CD4 | Biotin | GK1.5 | 1102020 | SONY | Rat | FC | Lineage Depletion (Hemato) |
| CD45 | BV786 | 30-F11 | 564225 | BD Biosciences | Rat | FC | Hematopoietic Cells |
| CD45 | Biotin | 30-F11 | 1115520 | SONY | Rat | FC | Lineage Depletion (Stroma) |
| CD45 | - | Polyclonal | AF114 | R&D Systems | Goat | IHC | Hematopoietic Cells |
| CD48 | APC | HM48-1 | 1117060 | SONY | Rat | FC | Hematopoietic Progenitors |
| CD51 | BV421 | RMV-7 | 740062 | BD Biosciences | Rat | FC | Mesenchymal Cells |
| CD71 | BV421 | RI7217 | 1169065 | SONY | Rat | FC | Erythroid Cells |
| CD71 | Biotin | RI7217 | 1169015 | SONY | Rat | FC | Lineage Depletion (Stroma) |
| CD71 | Biotin | RI7217 | 113803 | Biolegend | Rat | IHC | Erythroid Cells |
| CD8a | Biotin | 53-6.7 | 1103520 | SONY | Rat | FC | Lineage Depletion (Hemato) |
| Desmin | - | Polyclonal | ab8592 | abcam | Rabbit | IHC | Stellate Cells |
| Gp38 | APC-Cy7 | 8.1.1 | 1237090 | SONY | Rat | FC | Mesothelial Cells |
| Gp38 | - | 8.1.1 | NB600-1015 | Novus Biologicals | Hamster | IHC | Mesothelial Cells |
| Gr-1 | BUV395 | RB6-8C5 | 563849 | BD Biosciences | Rat | FC | Granulocytes |
| Gr-1 | Biotin | RB6-8C5 | 553125 | BD Biosciences | Rat | FC | Lineage Depletion (Hemato) |
| HNF4a | - | K9218 | MA1-199 | Invitrogen | Mouse | IHC | Hepatic Cells |
| HNF4a | - | EPR16885-99 | ab201460 | abcam | Rabbit | IHC | Hepatic Cells |
| LHX2 | - | Polyclonal | ABE1402 | Merck Millipore | Rabbit | IHC | Stellate cells |
| NG2 | PE | 1E6.4 | 130-100-468 | Milteny Biotec | Rat | FC | Mesenchymal Cells |
| NG2 | - | Polyclonal | AB5320 | Merck Millipore | Rabbit | IHC | Mesenchymal Cells |
| NK1.1 | Biotin | PK136 | 1143520 | SONY | Rat | FC | Lineage Depletion (Hemato) |
| Sca1 | BV510 | D7 | 565507 | BD Biosciences | Rat | FC | Hematopoietic Progenitors and Endothelial cells |
| Sca1 | BV711 | D7 | 1140655 | SONY | Rat | FC | Hematopoietic Progenitors and Endothelial cells |
| Ter119 | PE-Cy7 | TER-119 | 1181110 | SONY | Rat | FC | Erythroid Cells |
| Ter119 | Biotin | TER-119 | 1181020 | SONY | Rat | FC | Lineage Depletion (Hemato and Stroma) |
| Ter119 | Biotin | TER-119 | 116203 | Biolegend | Rat | IHC | Erythroid Cells |
| Streptavidin | BV785 | - | 2626245 | SONY | - | FC |  |
| Streptavidin | PE-Cy5 | - | 2626025 | SONY | - | FC |  |
| Streptavidin | AF488 | - | 405235 | Biolegend | - | IHC |  |
| Secondary Chicken anti-mouse | AF647 | - | A21463 | Invitrogen | - | IHC |  |
| Secondary Donkey anti-goat | AF488 | - | A11055 | Invitrogen | - | IHC |  |
| Secondary Donkey anti-goat | AF568 | - | A11057 | Invitrogen | - | IHC |  |
| Secondary Donkey anti-goat | AF647 | - | A21447 | Invitrogen | - | IHC |  |
| Secondary Donkey anti-mouse | AF488 | - | A21202 | Invitrogen | - | IHC |  |
| Secondary Donkey anti-mouse | AF568 | - | A10037 | Invitrogen | - | IHC |  |
| Secondary Donkey anti-rabbit | AF488 | - | A21206 | Invitrogen | - | IHC |  |
| Secondary Donkey anti-rabbit | AF568 | - | A10042 | Invitrogen | - | IHC |  |
| Secondary Donkey anti-rabbit | AF647 | - | A31573 | Invitrogen | - | IHC |  |
| Secondary Donkey anti-rat | AF488 | - | A21208 | Invitrogen | - | IHC |  |
| Secondary Donkey anti-rat | AF594 | - | A21209 | Invitrogen | - | IHC |  |
| Secondary Donkey anti-rat | AF647 | - | A48272 | Invitrogen | - | IHC |  |
| Secondary Donkey anti-sheep | AF568 | - | A21099 | Invitrogen | - | IHC |  |
| Secondary Donkey anti-sheep | AF647 | - | A21488 | Invitrogen | - | IHC |  |
| Secondary Rabbit anti-hamster | AF594 | - | 307-585-003 | Jackson ImmunoResearch | - | IHC |  |
| Anti-GFP | - | - | A11122 | Invitrogen | Rabbit | IHC |  |

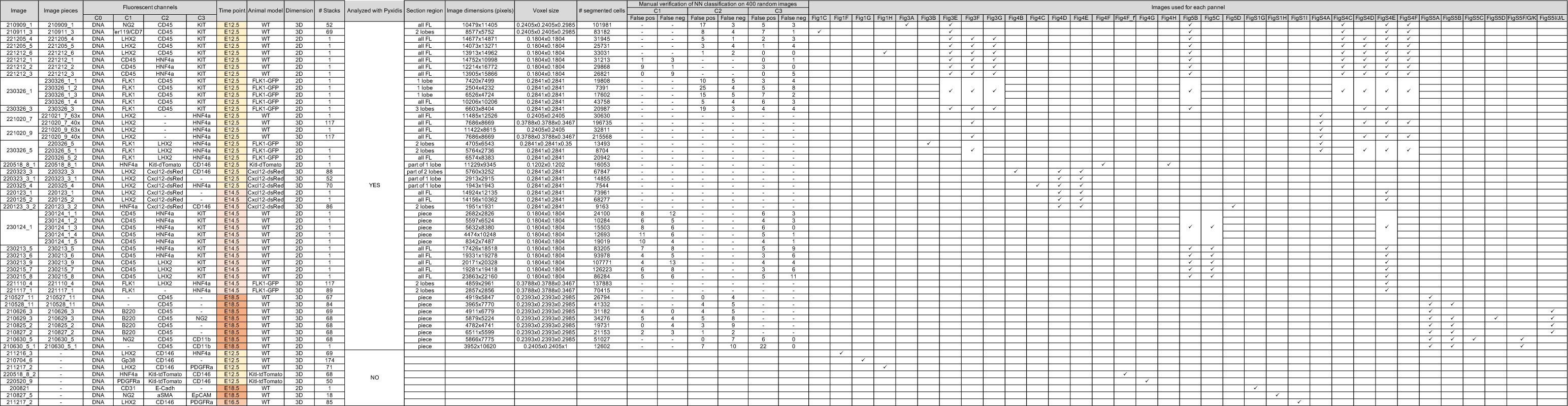

## Experimental Model and Subject Details

### Mice

C57BL/6J mice were purchased from Envigo. Flk1-GFP mice were a gift from David Hardy^(1)^. Fixed FLs from Cxcl12-dsRed^(2)^ mice were kindly provided by João Pedro Pereira. Fixed FLs from Kitl-tdTomato^(3)^ mice were kindly provided by Marella de Bruijn.

6 to 8-week-old mice or timed pregnant females were used. Timed pregnancies were generated after overnight mating, the following morning females with vaginal plugs were considered to be at E0.5. All animal manipulations were performed according to the ethics charter approved by the French Agriculture ministry and to the European Parliament Directive 2010/63/EU.

## Method Details

### Cell Suspension

For the isolation of non-hematopoietic populations, E12.5-E18.5 FLs were dissected under a binocular magnifying lens. FLs were recovered in Hanks’ balanced-salt solution (HBSS) supplemented with 1% fetal calf serum (FCS) (Gibco), cut in 1 mm^2^ pieces and incubated with 0,05 mg/ml Liberase TH (Roche) for 6-8 minutes at 37°C. Enzymatic treatment was stopped by addition of cold HBSS+1% FCS followed by centrifugation. FLs were depleted of hematopoietic cells (Ter119^+^CD45^+^Kit^+^CD71^+^) using MACS columns (Miltenyi Biotec) according to manufacturer instructions. Briefly, for MACS columns depletion, cells were incubated with biotinylated Ter119, CD45, Kit and CD71 antibodies for 20-30 min at 4°C, washed twice, incubated with anti-biotin microbeads (Miltenyi Biotec) for 20 min at 4°C and passed through a LS column (Miltenyi Biotec). The flow-through was collected and used for cell surface staining. Before staining, cell suspensions were filtered with a 100 μm cell strainer (BD).

For the isolation of hematopoietic cells, E12.5-E14.5 FLs were dissected under a binocular magnifying lens. FLs were recovered in HBSS+1% FCS (Gibco) and passed through a 26-gauge needle of a 1-ml syringe to obtain single-cell suspensions. Before staining, cell suspensions were filtered with a 100 μm cell strainer (BD).

For the isolation of BM non-hematopoietic populations, femurs and tibias were isolated from P7, P14, and P21 mice and surrounding tissues removed. Bones were flushed using a 26-gauge needle of a 1-ml syringe with HBSS+2% FCS (Gibco). BM clumps were re-suspended by gentling aspirating with a 19-gauge needle, filtered (100 μm) and kept on ice. Bones were crushed with a scissor in a way to maximize the internal bone surface area exposition and incubated with a digestion buffer (1mg/mL Collagenase VIII and X DNase I in HBSS+2% FCS) for 15 min at 37°C with rotation. Cell suspensions were collected, filtered (40 μm), and kept on ice. 3 rounds of digestion were performed, and cell suspensions collected from each were pooled together with the flushed ones. Enzymatic treatment was stopped by addition of cold HBSS+2% FCS followed by centrifugation. Cell suspensions were depleted of hematopoietic cells using MACS columns (Miltenyi Biotec) according to manufacturer instructions. Briefly, cells were incubated with biotinylated Ter119, CD45, CD11b and B220 antibodies for 20-30 min at 4°C, washed twice, incubated with anti-biotin microbeads (Miltenyi Biotec) for 20 min at 4 °C and passed through a LS column (Miltenyi Biotec). The flow-through was collected and used for cell surface staining. Before staining, cell suspensions were filtered with a 100 μm cell strainer (BD).

### Flow cytometry and cell sorting

MACS depleted non-hematopoietic cells were stained with the antibodies listed in Table S1 for 20 min at 4°C and analyzed on a spectral cytometer ID7000 (SONY) as described^(4)^ or sorted with a BD FACS Aria III (BD Biosciences) according to the guidelines for the use of flow cytometry and cell sorting^(5)^.

Hematopoietic cells were stained with the antibodies listed in Supp Table S1 for 20-30 min at 4°C and analyzed on a custom BD LSR Fortessa (BD Biosciences). For sorting, FL were depleted of mature cells using MACS columns (Miltenyi Biotec) according to manufacturer instructions. Briefly, cells were incubated with biotinylated Ter119, Gr-1, CD19, CD3, CD4, CD8, NK1.1, CD11c antibodies (listed in Supp Table S1) for 20-30 min at 4°C, washed, incubated with anti-biotin microbeads (Miltenyi Biotec) for 20 min at 4°C and passed through a LS column (Miltenyi Biotec). The flow-through was collected and used for cell surface staining with the antibodies listed in Table S1, for 20-30 min at 4°C. Stained cells were sorted with a BD FACSAria III (BD Biosciences) according to the guidelines for the use of flow cytometry and cell sorting^(5)^. Data were analyzed with FlowJo software (v.10.5.3, BD Biosciences).

### Flow Cytomety Analysis

Flow cytometry files were screened for abnormalities (flow rate, signal acquisitions, and dynamic range) using a quality control plugin in FlowJo, FlowAI^(6)^. Only ”GoodEvents” were considered for analysis. To perform dimensionality reduction of the flow cytometry data, UMAPs were performed in FlowJo, on gated single, live, non-hematopoietic cells. For each stromal population defined, the corresponding channel values for the UMAP parameters were saved as .csv and plotted using Jupyter Notebook (University of Southampton Institutional Repository).

## Imaging

### FL sectioning, immunostaining and optical clearing of thick sections

FLs from timed-pregnant females were collected directly into freshly prepared 2% PFA fixative buffer (Fischer Scientific) and kept at 4°C overnight under rotation. After washing in PBS and removing surrounding tissues, FLs were immediately sectioned or shortly stored at 4°C until sectioning. Following FLs embedding in 4% low-melting agarose (Sigma), 150 µm frontal sections were performed using a Leica VT1200S vibratome. Sections were permeabilized and blocked with PBS containing 0.05% Tween-20 (AppliChem GmbH) [or 0.5% Triton-X 100 for nuclear staining] and 10% donkey serum (Sigma-Aldrich) for 1-2 hr at RT. This solution was also used to dilute primary and secondary antibodies. Sections were incubated with primary antibodies for 2 hours at room temperature (RT) and overnight at 4°C, and secondary antibodies for 2h at RT, both in the dark and with rotation. Between incubations and after secondary antibody incubation, sections were washed repeatedly with PBS containing 0.05% Tween-20 and 3% NaCl (minimum 4x, 15 min). Sections were counterstained with DAPI (D9542, Sigma-Aldrich) for 2 hours at RT and kept overnight in RapiClear 1.52 (SunJin Lab) for optical clearing. Sections were mounted in RapiClear and imaged on a Leica TCS SP8 using a PL APO 63x /1.40 oil immersion lens or APO 40x /1.40 oil immersion lens. Single-stacks or multi-stacks tile scans were acquired at 200Hz or 400Hz, respectively, bidirectional mode, at 12-bits and 1024×1024 resolution. Multi-stacks were acquired with z spacing of 0.3 μm. Table S2 compiles all information regarding image acquisition.

## Image Analysis

### Image processing and segmentation

The first step to obtain these graphs consisted of segmenting the nuclei within each image, based on their DAPI signal, using CellPose^(7)^. The FL images acquired in this study ranged in size from 2682×2826 to 7686×8669×117 pixels (See Table S2), which corresponded to 60Mb up to 50Gb in memory needs. As a result, some were too large for analysis on graphics processing unit (GPU)s with 8GB RAM. These images were pre-processed to reduce memory consumption by dividing them into smaller tiles using a Python module named *Saucisson* before being segmented (Fig 2A-1). Tiles of 3000×3000 pixels were used for 2D images and 600×600×Z-stacks for 3D images (See Table S2 for number of z-stacks). These tiles size allow to fit data sizes to GPUs memory and parallel computations. These separate files were then segmented in parallel on a GPU cluster, using the Cellpose package (Fig. 2A-2). After segmentation, the masks from each of the individual tiles were stitched together again using Saucisson (Fig. 2A-3).

### Cell classification

As a result of the image segmentation pipeline, the position of each cell is known only by the segmented domain of its nucleus. The next step is to distinguish the cell phenotype, which is obtained from the expression levels of nuclear or membrane-bound proteins to which specific fluorescent antibodies can attach.

In order to classify cells based on staining in the nucleus (such as LHX2 and HNF4a), the average signal intensity, within the mask of the nucleus, is computed using *Griottes*^(8)^ and a threshold is applied, robustly distinguishing between positive and negative cells (Fig 2B-4). To classify cells based on membrane staining, it is necessary to analyze the fluorescence pattern around each nucleus. In this case measuring average fluorescence intensity inside or around the nucleus isn’t precise enough, since neighboring positive cells can cause a local high fluorescence signal, despite little or no expression of the relevant protein by the cell itself (Fig 2B-5). Instead, the cells were classified using a deep learning based Neural Network. An individual square image was generated for each detected nucleus, with dimensions between 18 to 22µm (80×80 to 120×120 pixels) and centered around the geometric center of the nucleus (See Fig.2B and S3 for examples). The gray-scale pattern of the fluorescence inside this box was then analyzed by using *SqueezeNet* networks^(9)^, using the transfer learning approach and loaded pre-trained from *PyTorch*^(10)^. Training and validation datasets were manually created (Fig 2B) for each staining, using data from different images. Training was performed using a batch size of 32. As a result, an accuracy larger than 95% was achieved on validation datasets for each membrane staining (Fig S3A-B), although some manual corrections in the classification were performed for images with lower staining contrast. A different network was trained for each marker in order to manage differences that appeared for the different membrane stainings. Manual verification of classification performances was done on 400 cells randomly selected for each membrane staining and each image. The number of false negative and positive cells is given in Table S2. The accuracy is larger than 96% for all verified set of cells.

### Removal and selection of specific regions

In FL sections, nucleated erythrocytes are frequently found in blood vessels. Due to a very specific auto fluorescent (AF) signal, intense cytoplasmatic signal in one or more channels, they can be misclassified for membrane staining (e.g., as CD45^+^ and/or KIT^+^). To correct this, we used the same neural network approach, previously described, to detect these cells, and cells identified as AF^+^ were excluded from analysis (Fig S3C-E). Some images also contain disrupted lobes and pieces of tissues coming from other organs. A specific interactive tool named *Coloriage* was created to graphically select regions in the segmented image. This tool is used to eliminate these undesired regions (Fig. 2C-6; Fig S3F).

**Figure 6.**
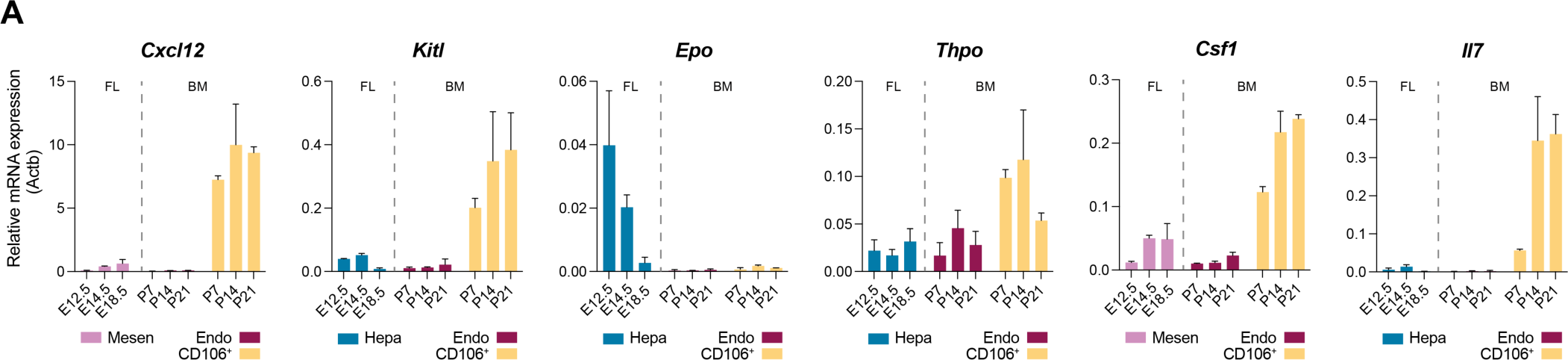
The fetal liver is a low cytokine environment. **(A)** Quantitative RT-PCR analysis of the expression levels of the hematopoietic cytokines *Cxcl12, Kitl, Epo, Thpo, Csf1* and *Il7* in mesenchymal (pink) or hepatic (blue) cells of E12.5, E14.5 and E18.5 FL and endothelial (dark red) or CD51^+^CD106^+^ (yellow) cells of P7, P14 and P21 BM. For each cytokine, only the FL population with highest expression levels is represented: Mesen (pink) for *Cxcl12* and *Csf1* and Hepa (blue) for *Kitl, Epo, Thpo* and *Il7* (n=3).

### Network construction

The outcome of this pipeline was to replace the pixel/voxel-scale description with a cell-scale representation of the complete image, either in 2D or in 3D. In this representation each cell was identified by its position and was characterized by its expression of the different nuclear or membrane markers (Fig. 2C). This representation was stored as a connected graph using the Griottes package^(8)^ (Fig. 2C-7). The connectivity network of the FLs was generated using *Griottes* from a data table containing the positions and properties of the nuclei cells using the ‘Delaunay’ construction rule. Further information on *Griottes* can be found in^(8)^. This network construction allows us to estimate the connectivity between any two cells and to build the statistics of proximity between specific cell types and stromal cells (Fig 2C).

The graph-based description provides a method to extract key information from large fluorescence images, including cell phenotype and position, and condensing it into a generic format. This condensation allows the data size to be reduced from several Gb (up to 50Gb for the analyzed images) to a few Mb per image, without loss of information. Moreover, by labeling specific nodes in the graph as being at the periphery, it is possible to calculate distances of cells from this periphery from independent images and to pool the information together, which in turn enables the identification of robust and recurring spatial patterns (as demonstrated in Figs. 3 and 5).

### Calculation of the cell enrichment in different regions

The relative enrichment or depletion of each cell type was calculated by comparing the relative abundance at any region with a random distribution of cell types. In particular, we focused on the enrichment near the periphery and the vessels in the FL. This procedure began by graphically selecting the specific structure to be studied, using *Coloriage*. The minimum Euclidean distance between each cell in the network and the cells attributed to the structure of interest was computed. Then the relative abundance of each cell type was binned as a function of distance from the structure.

In parallel the cell types were numerically shuffled on the network representation, while preserving the total frequency of the types in the tissue. The distance to the periphery was re-calculated for ten repetitions of the re-shuffling operation, thus providing a randomized position graph of the different cell types.

The ratio of the measured distribution to the randomized distribution provided a robust measure of the enrichment at different distances from the periphery and the vessels within the tissue. This allows us to distinguish relevant and statistically significant patterns that are not a consequence of random cell distribution.

## Code availability

The analysis on the FL network images was conducted on Jupyter Notebook. All the relevant notebooks are freely accessible on the project GitHub repository (https://github.com/BaroudLab/Pyxidis). Different packages have been custom-made for the analysis of the FL images, during pre-processing and data analysis. All these functionalities (*Saucisson*, *Coloriage*) have been incorporated in the main GitHub for easier use and installation. This repository contains a step-by-step tutorial to run the entire image analysis pipeline and data analysis previously described.

### Multiplex single-cell qPCR

Single cells were sorted directly into 96-well plates loaded with RT-STA Reaction mix (CellsDirect^TM^ One-Step qRTPCR Kit, Invitrogen, according to the manufacturer’s procedures) and 0.2x specific TaqMan® Assay mix (see Table S3 for the TaqMan® assays list) and were kept at −80°C at least overnight. For each subset analyzed, a control well with 20 cells was also sorted. Pre-amplified cDNA (20 cycles) was obtained according to the manufacturer’s note and was diluted 1:5 in TE buffer for qPCR. Multiplex qPCR was performed using the microfluidics Biomark HD system for 40 cycles (Fluidigm) as previously described^(11)^. The same TaqMan probes were used for both RT/pre-amp and qPCR. Only single cells for which at least 2 housekeeping genes could be detected before 20 cycles were included in the analysis.

### Gene expression by RT-PCR

Cells were sorted directly into lysis buffer and mRNA was extracted with a RNeasy Plus Micro Kit (Qiagen). After extraction, mRNA was reverse-transcribed into cDNA with PrimeScript RT Reagent Kit (Takara Bio), followed by quantitative PCR with Power SYBR Green PCR Master Mix (Applied Biosystems). Primers used can be found in Supplementary Table S4. qPCR reactions were performed on a Quantstudio3 thermocycler (Applied Biosystems), gene expression was normalized to that of b-actin and relative expression was calculated using the 2^-ΔCt^ method.

### Transwell migration assays

FL KIT^+^ and DP cells were prepared as for flow cytometry and cell sorting. For sorting of E14.5 cells, lineage depletion (Ter119, CD19, B220, Gr1, CD3, CD4, CD8a, NK1.1 and CD11c) was performed using magnetic LS columns (Miltenyi Biotec). Dual-chamber chemotaxis assays were performed using 24-well plates with 5-μm pore size inserts (Cat# PTMP24H48, Sigma), as previously described^(12)^. CXCL12a (Cat# 460-SD- 050, R&D Systems) was added to the lower chamber at defined concentrations, and 100 μl of a cell suspension (1 × 10^6^ cells/ml) of KIT^+^ or DP cells was placed in the upper chamber. Cells were allowed to migrate for 2h at 37°C. A known quantity of fluorescent beads was added to the lower chamber for normalization of migrated progenitors. Migrated cells were collected from the lower chamber and analyzed by FACS to enumerate migrated progenitors.

### Bioinformatic analysis of multiplex single-cell qPCR

Gene expression raw data (BioMarkTM, Fluidigm) of single cells was normalized with Gapdh and Actb. Heatmaps and hierarchical clustering were generated using R packages “pheatmap” and “Rphenograph”^(13)^.

### Quantification and Statistical Analysis

All results are shown as mean ± standard deviation (SD). Statistical significance was determined using two-way ANOVA, followed by Šídák’s multiple comparisons test where a P value of <0.05 was considered significant and a P value >0.05 was considered not significant.

---

**Table S3.** – Taqman Assays used for single-cell multiplex qPCR.

| Gene | Gene ID | Taqman Assay ID |
| --- | --- | --- |
| Acta2 | 11475 | Mm01546133_m1 |
| Afp | 11576 | Mm00431715_m1 |
| Alb | 11657 | Mm00802090_m1 |
| Alcam | 11658 | Mm00711623_m1 |
| Angpt1 | 11600 | Mm00456503_m1 |
| Angpt2 | 11601 | Mm00545822_m1 |
| Cdh1 | 12550 | Mm01247357_m1 |
| Cdh5 | 12562 | Mm00486938_m1 |
| Cebpa | 12606 | Mm00514283_s1 |
| Csf1 | 12977 | Mm00432686_m1 |
| Csf2 | 12981 | Mm01290062_m1 |
| Csf3 | 12985 | Mm00438334_m1 |
| Cspg4 | 121021 | Mm00507257_m1 |
| Cxcl12 | 20315 | Mm00445553_m1 |
| Des | 13346 | Mm00802455_m1 |
| Dlk1 | 13386 | Mm00494477_m1 |
| Efnb2 | 13642 | Mm00438670_m1 |
| Endoglin | 13805 | Mm00468252_m1 |
| Epcam | 17075 | Mm00493214_m1 |
| Ephb4 | 13846 | Mm01201157_m1 |
| Epo | 13856 | Mm01202755_m1 |
| Flt3l | 14256 | Mm00442801_m1 |
| Fn1 | 14268 | Mm01256744_m1 |
| Gpm6a | 234267 | Mm00463812_m1 |
| Hnf4a | 15378 | Mm01247712_m1 |
| Igf2 | 16002 | Mm00439564_m1 |
| Il33 | 77125 | Mm00505403_m1 |
| Il7 | 16196 | Mm00434291_m1 |
| Kdr | 16542 | Mm01222421_m1 |
| Kitl | 17311 | Mm00442972_m1 |
| Krt18 | 16668 | Mm01601704_g1 |
| Krt19 | 16669 | Mm00492980_m1 |
| Lepr | 16847 | Mm00440181_m1 |
| Lgr5 | 14160 | Mm00438890_m1 |
| Lhx2 | 16870 | Mm00839783_m1 |
| Ltb | 16994 | Mm00434774_g1 |
| Ly6a | 110454 | Mm00726565_s1 |
| mKi-67 | 17345 | Mm01278617_m1 |
| Msln | 56047 | Mm00450770_m1 |
| Msx2 | 17702 | Mm00442992_m1 |
| Nes | 18008 | Mm00450205_m1 |
| Ngfr | 18053 | Mm00446296_m1 |
| Onecut1 | 15379 | Mm00839394_m1 |
| Pdgfra | 18595 | Mm00440701_m1 |
| Pdgfrb | 18596 | Mm00435553_m1 |
| Pdpn | 14726 | Mm00494716_m1 |
| Prox1 | 19130 | Mm00435969_m1 |
| Runx1 | 12394 | Mm01213404_m1 |
| Sox4 | 20677 | Mm00486320_s1 |
| Sox9 | 20682 | Mm00448840_m1 |
| Spp1 | 20750 | Mm00436767_m1 |
| Tek | 21687 | Mm00443243_m1 |
| Thpo | 21832 | Mm00437040_m1 |
| Vcam1 | 22329 | Mm01320970_m1 |
| Vim | 22352 | Mm01333430_m1 |
| Wt1 | 22431 | Mm01337048_m1 |
| Actb | 11461 | Mm01205647_g1 |
| Gapdh | 14433 | Mm99999915_g1 |
| Hprt | 15452 | Mm03024075_m1 |

**Table S4.**
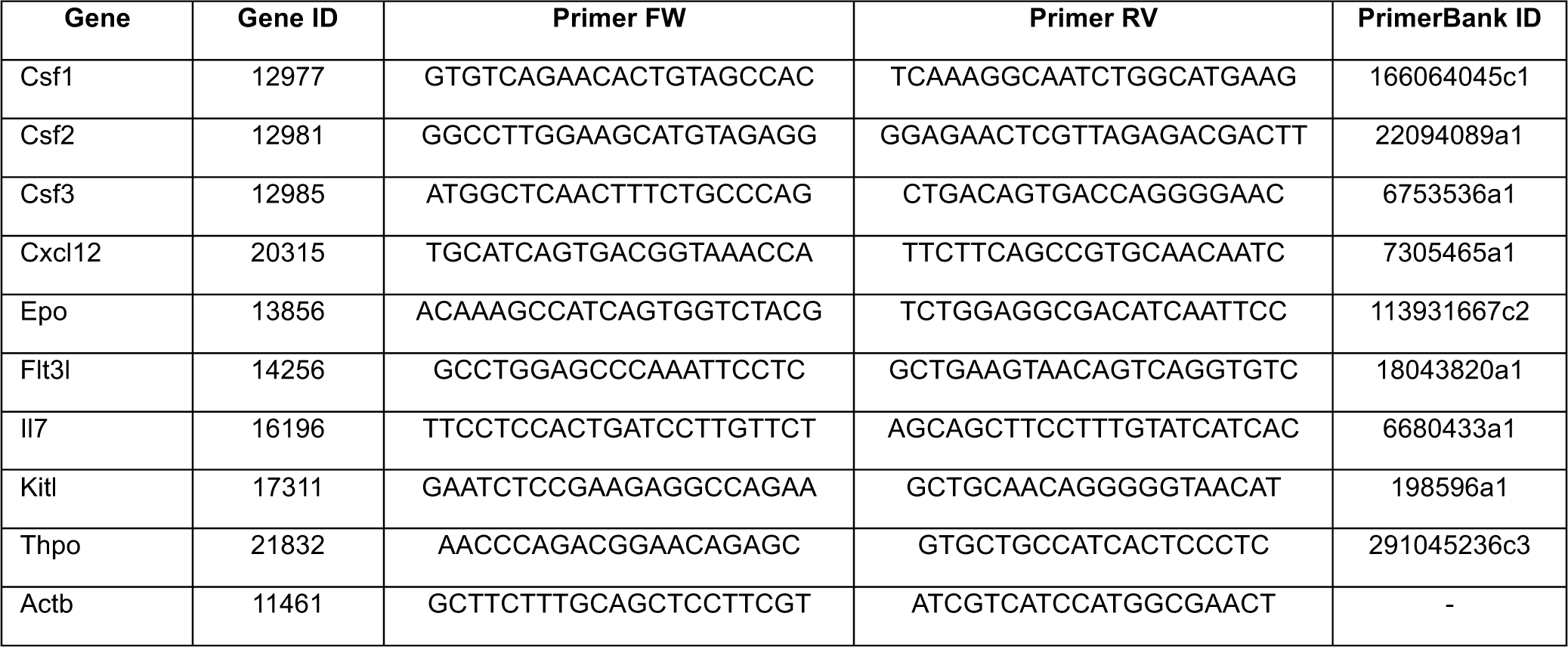
- Primers used for RT-PCR.

